# Activity-dependent induction of astrocytic Slc22a3 regulates sensory processing through histone serotonylation

**DOI:** 10.1101/2023.02.24.529904

**Authors:** Debosmita Sardar, Yi-Ting Cheng, Junsung Woo, Dong-Joo Choi, Zhung-Fu Lee, Wookbong Kwon, Hsiao-Chi Chen, Brittney Lozzi, Alexis Cervantes, Kavitha Rajendran, Teng-Wei Huang, Antrix Jain, Benjamin Arenkiel, Ian Maze, Benjamin Deneen

**Affiliations:** Center for Cell and Gene Therapy, Baylor College of Medicine, Houston TX; Center for Cancer Neuroscience, Baylor College of Medicine, Houston TX; Program in Developmental Biology, Baylor College of Medicine, Houston TX; Program in Development, Disease Models, and Therapeutics, Baylor College of Medicine, Houston TX; The Integrative Molecular and Biomedical Sciences Graduate Program, Baylor College of Medicine, Houston TX; Genetics and Genomics Graduate Program, Baylor College of Medicine, Houston TX; Mass Spectrometry Proteomics Core, Baylor College of Medicine, Houston TX; Department of Molecular and Human Genetics, Baylor College of Medicine, Houston TX; Neurological Research Institute, Texas Children’s Hospital, Houston TX; Nash Family Department of Neuroscience, Friedman Brain Institute, Icahn School of Medicine at Mount Sinai, New York NY; Department of Pharmacological Sciences, Icahn School of Medicine at Mount Sinai, New York NY; Howard Hughes Medical Institute, Icahn School of Medicine at Mount Sinai, New York NY 10029; Department of Neurosurgery, Baylor College of Medicine, Houston TX 77030

**Author notes:** These authors have equal contribution (Yi-Ting Cheng, Junsung Woo).

## Abstract

Neuronal activity drives global alterations in gene expression within neurons, yet how it directs transcriptional and epigenomic changes in neighboring astrocytes in functioning circuits is unknown. Here we show that neuronal activity induces widespread transcriptional upregulation and downregulation in astrocytes, highlighted by the identification of a neuromodulator transporter Slc22a3 as an activity-inducible astrocyte gene regulating sensory processing in the olfactory bulb. Loss of astrocytic Slc22a3 reduces serotonin levels in astrocytes, leading to alterations in histone serotonylation. Inhibition of histone serotonylation in astrocytes reduces expression of GABA biosynthetic genes and GABA release, culminating in olfactory deficits. Our study reveals that neuronal activity orchestrates transcriptional and epigenomic responses in astrocytes, while illustrating new mechanisms for how astrocytes process neuromodulatory input to gate neurotransmitter release for sensory processing.

## Introduction

Astrocytes are intimately associated with neuronal activity and neurotransmission, participating in a host of essential roles that facilitate synaptic function (*1–5*). A series of prior studies have shown that cell intrinsic activation of Gq- or Gi-signaling and associated Ca^2+^ activity in astrocytes can influence a multitude of behavioral outputs, providing key evidence for astrocytic modulation of circuit function (*6–10*). However, in native behavioral states astrocyte activation results from cell extrinsic sources, usually in response to a neuronal stimulus, and therefore typically follows neuronal activation (*11*). In neurons, heightened activity induces expression of ‘immediate early genes’, which are predominately transcription factors that modify gene expression programs and activity-dependent epigenomic states, ultimately regulating circuit activity, plasticity, and associated behavioral outputs (*12*). A series of recent studies have identified astrocytic transcription factors that regulate region specific-circuits in the adult brain (*13, 14*), indicating that transcriptional states in astrocytes can influence neuronal activity. Evidence for the reciprocal relationship can be found in the developing cortex (*15*) and *in vitro* (*16, 17*), where alterations in neuronal activity influences expression of synaptogenic genes in differentiating astrocytes. Single-cell transcriptomic studies have shown non-neuronal cells undergo transcriptional changes in response to neuronal activity driven by visual stimuli (*18*). However, whether heightened neuronal activity induces an analogous ‘immediate early gene-like’ response in mature astrocytes and how this sculpts astrocytic transcriptional and epigenomic responses to regulate circuit function remains unclear.

A central component of astrocyte-neuron communication is synaptic neurotransmitter signaling, as astrocytes express a host of receptors and transporters for both glutamate and GABA, activation of which can elicit Ca2^+^ dependent responses that modify circuit activity (*5, 19*). Accordingly, maintenance of extracellular glutamate levels via astrocytic glutamate transporters are required for drug-seeking behaviors, circadian rhythms, and avoidance behaviors, among others (*20–23*). Similarly, disruption of GABA signaling in astrocytes leads to repetitive behaviors, impaired motor function, and deficits in learning and memory (*7, 24–26*). Another set of neuroactive compounds mediating astrocyte-neuron interactions is volume transmission-based neuromodulators, as astrocytes express a host of neuromodulator receptors and respond to noradrenaline, acetylcholine, and serotonin with intracellular Ca2^+^ elevations (*16*). Corresponding manipulations of cholinergic inputs to hippocampal astrocytes influences sleep states, while noradrenaline primes cortical astrocyte responses to local neuronal activity during locomotion, and dopamine promotes astrocyte-mediated depression of excitatory synapses, which influences drug-induced locomotor activity (*27–29*). Despite direct roles in gating responses to neuromodulators, the signaling mechanisms utilized by astrocytes to respond- and process-neuromodulatory cues remain unclear. Furthermore, how astrocytes coordinate the integration of neuromodulator (i.e., volume-based) with neurotransmitter (i.e. synaptic) signaling to regulate circuit function is unclear.

Epigenomic regulation of gene expression plays a critical role in encoding information in the central nervous system (CNS), where alterations in synaptic activity induce epigenetic modifications that influence behavioral outcomes (*12*). For example, alterations in histone acetylation are directly linked to memory storage and synaptic plasticity in hippocampal neurons after learning (*30*). A series of recent studies have shown that neuromodulators can be directly incorporated into histones and serve as a new form of epigenetic gene regulation in the CNS (*31, 32*). Among neuromodulators, serotonin and dopamine can be incorporated into histones, with histone serotonylation enabling permissive gene expression states in serotonergic neurons (*31*) and histone dopaminylation regulating cocaine seeking behaviors in rodents (*32*). Nevertheless, this remains a relatively nascent field, with several aspects of this biology remaining unclear, including the extent and significance of these histone modifications throughout the brain and the fundamental mechanisms of how neuromodulators are transported to the nucleus and added to histones. Furthermore, how these and other epigenomic mechanisms are utilized by astrocytes to regulate gene expression, circuit function, and associated behavioral responses are unknown.

In this study, we examined how neuronal activation impacts transcriptional responses in astrocytes, identifying robust activity-dependent transcriptomic changes. These studies uncovered the neuromodulator transporter Slc22a3 in olfactory bulb (OB) astrocytes as a key regulator of circuit function and associated olfactory responses. Mechanistically, Slc22a3 transporter regulates transport of serotonin into astrocytes and coordinates histone serotonylation in astrocytes to control the expression of GABA-associated genes and olfactory responses. Taken together, these results identify novel transcriptional and epigenetic mechanisms, while revealing how astrocytes integrate neuromodulator signaling to regulate sensory processing of olfaction.

### Neuronal activity directs transcriptional responses in astrocytes

To understand how neuronal activation influences transcriptional responses in astrocytes we employed chemogenetic approaches to activate neurons in Aldh1l1-GFP reporter mice (**Fig. 1A-B**). We used postnatal intraventricular injection of viral expression vector (pAAV) for hM3Dq DREADD (Gq-DREADD) under the pan-neuronal synapsin promoter, which allowed widespread neuronal Gq-DREADD expression throughout the brain. When mice reached early adulthood (∼8 weeks) we activated neuronal Gq signaling via intraperitoneal injection of clozapine-N-oxide (CNO) (0.3 mg/kg) or saline (represented as Gq-Saline or Gq-CNO herein), and harvested brains 30 minutes after treatment (**Fig. 1C**). We confirmed enhanced neuronal activity in Gq-CNO treated samples using electrophysiology in brain slice preparations, coupled with expression of neuronal immediate early gene marker c-Fos (**Supp Fig. S1A-C**). To identify neuronal activity-dependent gene expression changes in astrocytes, we utilized the presence of distinct reporters in Gq-DREADD expressing neurons (mCherry) and astrocytes (GFP) to isolate these two cell types from the cortex (CX), hippocampus (HP), and olfactory bulb (OB) (**Fig. 1D**) by fluorescent activated cell sorting (FACS) and performed RNA-sequencing (RNA-Seq) (**Supp Fig. S1D**). Analysis of Gq-CNO vs. Gq-Saline cohorts from each region revealed widespread changes in gene expression in astrocytes, identifying 267, 300 and 256 astrocyte-specific differentially expressed genes (DEGs) in the CX, HP, and OB, respectively (**Fig.1E-F**). These DEGs were specific to astrocytes, demonstrating that immediate early responses in neurons are distinct from those in astrocytes (**Supp Fig. S2A-C**). Furthermore, gene ontology (GO) analyses revealed distinct terms associated with astrocytes and neurons across the CX, HP and OB: top astrocytic GO terms involved protein phosphorylation, calcium ion transport and serotonin transport, in the CX, HP, and OB, respectively (**Supp Fig. S2D-F**; **Supp Table S1**). To control for possible transcriptional artifacts due to CNO treatment, we treated Aldh1l1-GFP mice with 0.3 mg/kg CNO (represented as CNO-control herein), harvested their OBs after 30 minutes, and FACS isolated GFP-expressing astrocytes, which was followed by RNA-Seq (**Supp Fig. S3A**). Comparing the RNA-Seq profiles between the untreated OB astrocytes and CNO-controls identified widespread changes in gene expression (**Supp Fig. S3B**). However, these CNO-induced changes do not correspond to any of the GO groups we identified in our activity-based profiling experiments (**Supp Fig. S3C**). Moreover, comparison of the activity-induced DEGs with the CNO-associated DEG’s revealed miniscule overlap, as amongst the 256 DEGs induced in OB astrocytes by neuronal activity, only 7 of these also demonstrated altered expression in the CNO-treated (**Supp Fig. S3D-E**), reinforcing the specificity of our activity-induced set of DEGs. Together, these data indicate that acute neuronal activation stimulates distinct widespread and region-specific changes in astrocytic gene expression throughout the brain.

**Figure 1.**
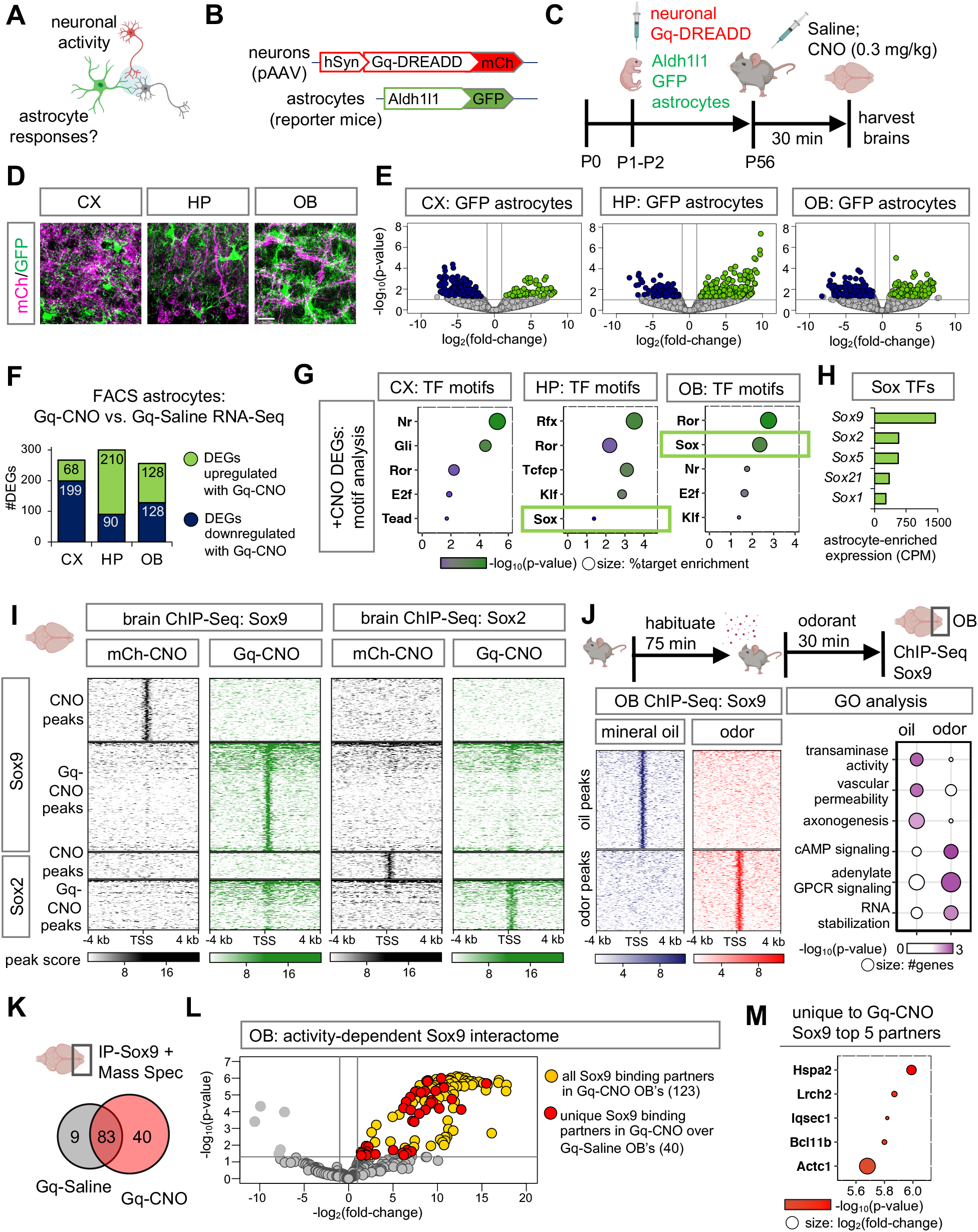
Neuronal activity directs Sox9-regulated transcriptional responses in astrocytes. (**A-C**) Schematic illustrating chemogenetic neuronal activation experimental design. (**D**) Sections showing distinct labeling of astrocytes (GFP) and Gq-DREADD neurons (mCh) in CX, cortex; HP, hippocampus; OB, olfactory bulb. Scale bar: 25 μm. (**E**) Volcano plots depicting RNA-Seq from GFP astrocytes comparing Gq-CNO vs. Gq-Saline. (**F**) Number of differentially expressed genes (DEGs) that are upregulated or downregulated in GFP astrocytes in Gq-CNO vs. Gq-Saline (n= 3/cohort, *p*< 0.05, log_2_fold-change 1). (**G**) Significant transcription factor (TF) motifs (*p*< 0.05) enriched in these DEGs and exhibiting astrocyte-specific expression. (**H**) Average transcript expression of Sox family TF’s in GFP astrocytes (CPM: counts per million). (**I**) Comparison showing heatmaps of ChIP-Sox9 and ChIP-Sox2 at 4 kb from peak center in Gq-CNO vs. mCh-CNO control (n= 3/cohort). (**J**) Schematic for odor evoked neuronal activation in the OB. Left panel: heatmaps of ChIP-Sox9 at 4 kb from peak center in mineral oil vs. odor exposed mice. Right panel: enriched gene ontology terms associated with these peaks (n= 6/cohort). (**K**) Sox9 binding partners in OB’s from Gq-CNO vs. Gq-Saline. (**L**) Volcano plot depicting IP-MS (IP-Mass Spectrometry) data of Sox9 interactome in Gq-CNO. Fold change was calculated over control lysates incubated with beads only without antibody. Sox9 binding partners unique to Gq-CNO vs. Gq-Saline are highlighted in red (n= 9–12/cohort, *p*< 0.05, log_2_fold-change >1). (**M**) Top 5 Sox9 interactors unique to Gq-CNO (*p*< 0.0001, log_2_fold-change >8.5). Color code represent p-values (x-axis), and size represent log_2_fold-change with larger circles denoting greater binding affinity.

To dissect the transcriptional mechanisms that underly these robust changes in astrocyte gene expression, we used transcription factor (TF) motif analysis on the astrocyte DEGs to nominate candidate TFs responsible for these changes in gene expression. Filtering for TFs that also exhibit enriched expression in astrocytes (**Supp Fig. S4A**) identified a cohort of TFs with motifs associated with activity-dependent DEGs in astrocytes, including the Sox-family of TFs in both the HP and OB (**Fig. 1G**). The Sox family has multiple members exhibiting elevated expression in astrocytes, with Sox9 and Sox2 being the most significant (**Fig. 1H**). Sox9 has been shown to be critical for olfactory circuit processing (*13*), while Sox2 has been shown to play roles in responses to brain injury (*33*). We next evaluated Sox9 and Sox2 protein levels after neuronal activation and observed no significant changes in protein expression measured by both immunostaining (**Supp Fig. S4B**) and immunoblot blot (**Supp Fig. S4C-D**). Therefore, we next asked if these Sox TFs undergo changes in their transcriptional activity after neuronal activation. Towards this, we performed chromatin immunoprecipitation coupled with next generation sequencing (ChIP-Seq) of Sox9 and Sox2 in CNO treated Gq-DREADD (Gq-CNO) expressing mice in comparison to empty pAAV-mCh expressing mice (represented as mCh-CNO herein), harvesting brains 30 minutes after CNO treatment. Although both Sox9 and Sox2 are equally immunoprecipitated (IP) in Gq-CNO and mCh-CNO control lysates (**Supp Fig. S5A**), we found that both of these Sox TFs exhibit >1.5-fold increase in DNA binding capacity in Gq-CNO brains compared to mCh-CNO controls (**Fig. 1I; Supp Fig. S5B-C**).

Furthermore, differential ChIP-Seq peaks analysis between Gq-CNO treatment vs. controls revealed 295 Sox9-specific peaks, 139 Sox2-specific peaks and 117 Sox9 and Sox2 shared peaks representing DNA binding activities of these Sox TFs is enhanced after neuronal activation (**Supp Fig. S5D**). GO analysis revealed activity-dependent peaks were associated with dendrite development (Sox9-specific), proteoglycans (Sox2-specific), while Sox9 and Sox2 shared peaks were enriched in metabolic and mRNA-processing (**Supp Fig. S5E**).

The forgoing studies were conducted after an artificial stimulation of neurons, leading us to examine whether such drastic changes in TF DNA binding would occur after native stimulation. Because Sox9 exhibited greater DNA binding capacity, and we recently demonstrated that astrocytic Sox9 is critical for olfactory sensory circuit function in the OB (*13*), we next examined whether olfactory sensory input driven neuronal activation influences Sox9 transcriptional activity. Importantly, in the OB, Sox9 expression is restricted to astrocytes (>96% Aldh1l1-GFP+ cells) and has miniscule expression in the oligodendroglial lineage (2.79% Olig2+ cells) (**Supp Fig. S6A-B**). To examine Sox9 transcriptional activity after native olfactory driven neuronal stimuli, we next performed Sox9 ChIP-Seq on OBs from mice sacrificed after exposure to odorant isoamyl acetate or mineral oil control for 30 minutes, revealing largely distinct Sox9 peaks and their associated GO’s (**Supp Figure S6C**). While Sox9 peaks in mineral oil exposed OBs were enriched in transaminase activity GO terms, peaks in odor exposed OBs were enriched in cAMP signaling GO terms (**Fig. 1J**). To assess if these changes in Sox9 DNA binding capacity after neuronal activation are due to changes in Sox9 binding partners, we next performed Sox9-IP coupled with mass spectrometry and observed a >4-fold increase in Sox9 interactions after neuronal activation (**Fig. 1K**; **Supp Table S2**), identifying 40 proteins that specifically interact with Sox9 in the Gq-CNO treated OBs in comparison to Gq-Saline control (**Fig. 1L**). Interestingly, we observed overlap between the top activity-specific Sox9 interacting proteins (**Fig. 1M**) and TFs identified from motif analysis of activity-specific ChIP-Sox9 peaks in the OB (**Supp Fig. S6D**). Collectively, these results demonstrate that neuronal activity induces robust transcriptional changes in astrocytes and regulates the function of key astrocyte TFs.

### Slc22a3 is an activity-dependent gene target of Sox9 in OB astrocytes

To identify astrocyte genes that are induced after neuronal activity, we used Sox9 and the OB as a model system. We compared the activity-dependent astrocyte-specific DEGs in the OB (**Fig. 1E**), with existing RNA-Seq data from OB astrocytes from Sox9 conditional knockout mouse (Sox9-cKO) (*13*), where Sox9 is deleted specifically from OB astrocytes. We identified 62 genes that are activity-dependent and Sox9 regulated, representing 24.2% of activity-dependent DEGs in OB astrocytes (**Supp Fig. S7A**). Among these, the 34 genes demonstrating decreased expression in the Sox9-cKO (**Fig. 2A**; **Supp Fig. S7B**), were further filtered based on Sox9 motif binding score at their promoters (**Fig. 2B**). Next, we performed a ChIP-PCR screen. In this screen, we immunoprecipitated DNA bound to Sox9, and performed PCR (**Supp Fig. S7C**) using primers spanning Sox9 motifs at promoter regions of candidate genes. Among the candidates demonstrating Sox9 promoter occupancy in the OB specifically after Gq-based neuronal stimulation was *Slc22a3* (**Fig. 2C-D**; **Supp Fig. S7D**). To validate Sox9-regulation of Slc22a3, we evaluated its protein expression in mice where Sox9 was specifically knocked out in adult astrocytes (Sox9-cKO; **Supp Fig. S8A**) (*13*), finding a drastic reduction of its expression in astrocytes from the OB (**Fig. 2E-F**), but not CX or HP (**Supp Fig. S8B-E**).

**Figure 2.**
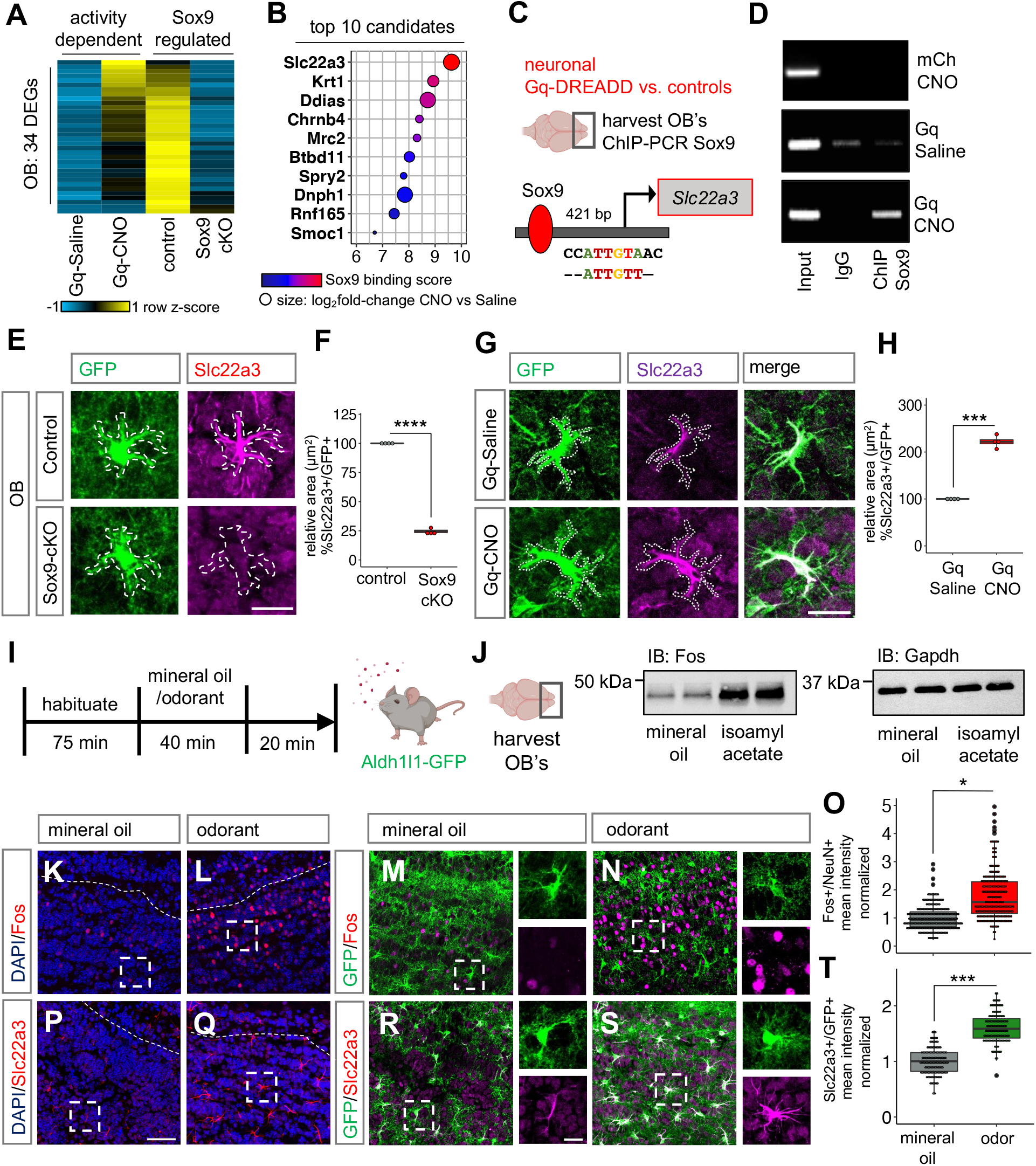
Neuronal activation induces Slc22a3 in OB astrocytes. (**A-B**) Heatmap depicting the 34 neuronal activity-dependent and Sox9 regulated DEGs (n= 3, *p*< 0.05), and the top 10 candidates filtered by Sox9 motif binding score 1000 bp from transcription start site of candidates. (**C-D**) ChIP-PCR screen identifying activity-dependent Sox9 regulation at *Slc22a3* promoter (n= 4-6 OB). (**E-H**) Slc22a3 expression in control vs. Sox9-cKO OB astrocytes and in Gq-Saline vs. Gq-CNO; and box plots depicting area covered by Slc22a3 in GFP astrocytes (average of 116-122 cells/cohort, \*\*\*\**p*= 1.93E-08; \*\*\**p*= 0.00013, unpaired Student’s two-tailed t-test on n= 4 mice/cohort); Scale bar: 20 μm. (**I**) Schematic illustrating odor-evoked neuronal activation experimental design. (**J**) Immunoblots of Fos and loading control from OB lysates (n= 4/cohort). (**K-O**) Immunostaining of Fos in Aldh1l1-GFP mice and quantification of mean fluorescence intensity in NeuN+ neurons (244-250 cells/cohort, \**p*= 0.0285, Wilcoxon rank sum test on n= 4 mice/cohort). (**S-T**) Immunostaining of Slc22a3 in Aldh1l1-GFP mice and quantification of mean fluorescence intensity in GFP+ astrocytes (130-131 cells/cohort, \*\*\**p*= 0.00039, unpaired Student’s two-tailed t-test on n= 4 mice/cohort). Scale bar: 50 μm; inset: 10 μm. Dashed line represents boundary of granule cell layer.

Next, we examined whether astrocytic Slc22a3 is regulated by neuronal activity by evaluating its protein and transcript expression in our Gq-DREADD based paradigm, where we found that its expression is upregulated in Gq-CNO treated OB astrocytes in comparison to Gq-Saline and CNO-control OBs (**Fig 2G-H; Supp Fig. S9A-E**). Next, we sought to determine whether neuronal activation from olfactory sensory input promotes Slc22a3 expression in the OB. We exposed habituated Aldh1l1-GFP mice to mineral oil or the odorant isoamyl acetate. Exposure was maintained for 40 minutes and after removal of oil or odorant, mice were kept in the chamber for an additional 20 minutes to allow protein turnover (**Fig. 2I**). To confirm increased neuronal activity in this paradigm, we evaluated Fos expression, a marker of neuronal activity, revealing increased protein expression in lysates obtained from the OB (**Fig. 2J; Supp Fig. S10A**) and in neurons of odor exposed OBs compared to mineral oil control, but not in astrocytes (**Fig. 2K-O**; **Supp Fig. S10B**). Next, we assessed Slc22a3, observing a significant increase in its astrocytic expression in odor exposed mice compared to mineral oil and non-habituated controls (**Fig. 2P-T; Supp Fig S10C-E**). These analyses were performed in the granular cell layer of the OB given both Fos and Slc22a3 enrichment in this sub-region of the OB. Moreover, we observed a small increase in the number of astrocytes that express Slc22a3 (**Supp Fig. S10F**) and increased levels of Slc22a3 expression was observed only in astrocytes and not in neurons as evaluated by co-labeling with NeuN (**Supp Fig. S10G**). These results indicate that astrocytic Slc22a3 is regulated by neuronal activation in the OB, and also highlights a unique set of astrocyte genes regulated by neuronal activity and Sox9.

### Astrocytic Slc22a3 is required for olfactory sensory processing

Slc22a3 belongs to the solute carrier family of membrane transport proteins and is involved in both release and uptake of the catecholamine family of chemicals, which include neuromodulators serotonin, dopamine and noradrenaline (*34–38*). While prior studies reported expression of Slc22a3 in astrocytes (*36, 37*) (**Supp Fig. S11A-B**), its contributions to astrocyte function and associated brain circuits are unknown. Given that odor input induces upregulation of astrocytic Slc22a3, we next determined whether Slc22a3 contributes to OB astrocyte function and olfactory circuits. To achieve selective knockout of Slc22a3 we generated a transgenic mouse line containing a floxed Slc22a3 allele (Slc22a3-FF), in combination with OB injection of pAAV containing Cre that is driven by the astrocyte specific Gfap promoter (**Supp Fig. S11C-D**). Injections occurred at P60, mice were harvested at P90, which enabled region- and cell type-specific deletion of Slc22a3 in OB astrocytes (**Supp Fig. S11E-F**), but not in neurons (**Supp Fig. S11G-I**). To quantify Slc22a3 knockout efficiency in astrocytes, we generated *Slc22a3-FF; Aldh1l1-GFP*, a double transgenic mouse line obtained by crossing Slc22a3FF and Aldh1l1-GFP mice. For control experiments, we injected Gfap-mCherry and for knockout experiments, we injected Gfap-Cre-RFP viral vectors; as before mice were injected at P60 and harvested at P90. (**Fig. 3A**). Quantification of Slc22a3 knockout efficiency across all Aldh1l1-GFP OB astrocytes revealed a significant reduction in Slc22a3 expression in astrocytes (**Fig. 3B-C**; **Supp Fig. S12A-D**), but not in neurons (**Fig. 3D-E**). Finally, we did not observe any changes in the overall numbers of astrocytes and neurons after Slc22a3 deletion in the OB (**Supp Fig. S12E-F**).

**Figure 3.**
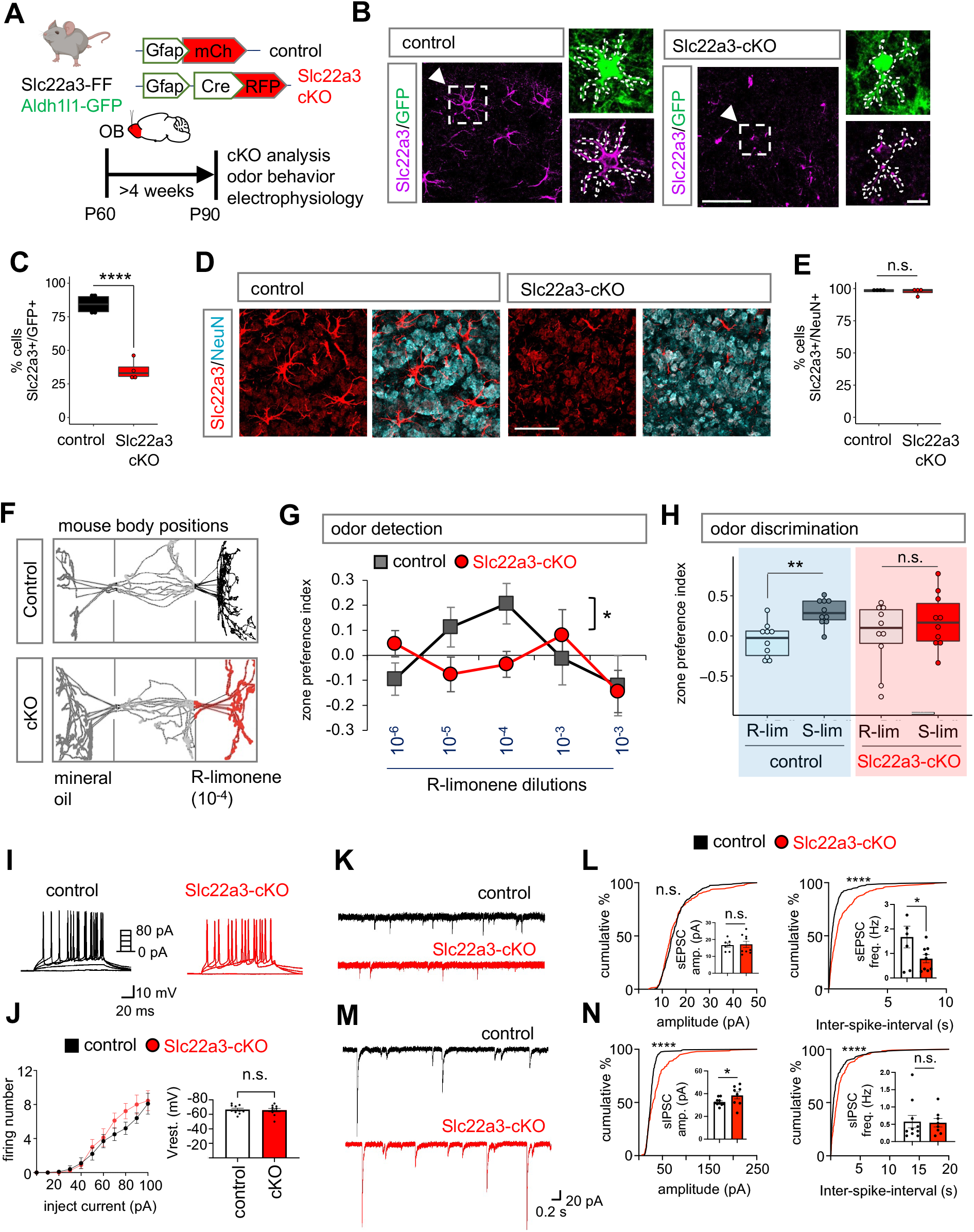
Astrocytic Slc22a3 regulates olfactory circuit function. (**A**) Schematic illustrating viral vectors used for Slc22a3 conditional deletion from OB astrocytes. (**B-E**) Representative immunostaining and quantification of Slc22a3 in GFP astrocytes (average of 83-93 cells/cohort, \*\*\*\**p*= 9.59E-05, unpaired Student’s two-tailed t-test on n= 4 mice/cohort); and in NeuN neurons (average of 350-363 cells/cohort, *p*= 0.5828, unpaired Student’s two-tailed t-test on n= 4 mice/cohort). Scale bar: 50 μm, inset: 10 μm. (**F**) Schematic illustrating live mouse tracking in three-chamber assay for odor detection. Top represent a control mouse exploring R-limonene (R-lim) at dilution of 10^−4^, while bottom represent Slc22a3-cKO mouse showing no preference for same. (**G**) Quantification of odor detection in control and Slc22a3-cKO mice (n= 10/cohort, \**p*= 0.0211; two-way repeated measures ANOVA with Sidak multiple comparison). (**H**) Quantification of odor discrimination between R-lim and S-limonene (S-lim) from the same cohorts of mice (n= 10/cohort, **p= 0.0052; two-way repeated measures ANOVA with Sidak multiple comparisons). (**I-J**) Whole-cell patch clamp electrophysiology of granule cells firing number from stepped current injections (n= 3, 10 cells/cohort, *p*= 0.0578, two-way ANOVA with Sidak’s multiple comparison correction). (**K-L**) Representative traces and summary data of amplitude and frequency from sEPSC recordings (7-9 cells/cohort, sEPSC amplitude *p=* 0.7546; sEPSC frequency \**p*= 0.0473, unpaired Student’s two-tailed t-test on n= 3 mice/cohort; \*\*\*\**p<* 0.0001 K-S test). (**M-N**) Representative traces and quantification of amplitude and frequency from sIPSC recordings (8-10 cells/cohort, sIPSC amplitude \**p*= 0.0164, sIPSC frequency *p*= 0.8095, unpaired Student’s two-tailed t-test on n= 3 mice/cohort \*\*\*\**p<* 0.0001 K-S test). All recordings in (L, N) are in granule cells and data is presented as mean ± SEM.

To determine how loss of Slc22a3 in OB astrocytes affects olfactory behaviors, we generated Slc22a3-cKO and controls as shown in Figure 3A. Next, we performed behavioral assays on these mice to evaluate potential alterations in odor behaviors (**Supp Figure S13A**). Odor detection was performed by olfactory habituation/dishabituation in a 3-chamber place preference assay, where mice are exposed to a novel odorant (R)-limonene in an increasing concentration series (**Supp Fig. S13B**). Odor detection was calculated based on the time spent investigating the odorant containing chamber in comparison to mineral oil chamber and represented as zone preference index. We found that Slc22a3-cKO required a higher concentration of odorant to exhibit preference for R-limonene, compared to controls, indicating impaired olfactory detection (**Fig. 3F-G**). Next, we evaluated odor discrimination, using a similar paradigm, where we compared the preference for structurally similar odorants (**Supp Fig. S13B**). While control mice spent more time in the new odorant containing chamber, Slc22a3-cKO mice showed no significant difference in preference, suggesting Slc22a3-cKO mice are deficient in their ability to discriminate between chemically similar, but distinct odorants (**Fig. 3H**). These defects in olfactory behaviors, which are coupled with enriched expression of c-Fos and Slc22a3 in the granular cell layer, led us to examine whether the electrophysiological properties of OB neurons are impacted by astrocytic knockout of Slc22a3. We performed whole-cell recordings from granule cells in the OB and found that excitability of granular cell layer neurons was unaffected (**Fig. 3I-J**).

Examination of their synaptic properties revealed a significant decrease in sEPSC frequency (**Fig. 3K-L**), coupled with an increase in sIPSC amplitude (**Fig. 3M-N**) in Slc22a3-cKO mice. Taken together, these results indicate that astrocytic Slc22a3 contributes to sensory processing in the OB through regulation of synaptic activity in granular cell neurons.

### Slc22a3 regulates astrocytic Ca^2+^ responses to neuromodulators

The forgoing results indicate that loss of astrocytic Slc22a3 disrupts olfactory function, leading us to examine how its loss impacts core features of astrocytes in the OB. Towards this, we performed RNA-Seq on FACS isolated astrocytes from control and Slc22a3-cKO OBs to identify molecular features impacting core astrocyte properties after loss of Slc22a3 (**Fig. 4A**). We found that Slc22a3 deletion (**Supp Fig. S13C)** dramatically affected astrocyte transcriptomics as revealed by 565 upregulated and 1049 downregulated genes that were differentially expressed in Slc22a3-cKO RFP astrocytes in comparison to empty mCh injected control astrocytes (**Fig. 4B, Supp Table S4**). GO analysis of these DEG’s revealed synaptic transmission as the most impacted category, in addition to cell-cell adhesion, interferon response, neurotransmitter transport, potassium ion transport and calcium ion binding. Constituent genes of all these categories were mostly downregulated, with the exception of interferon response genes, which were upregulated (**Fig. 4C-D**). Given that synaptic transmission is affected by astrocyte-neuron communication, which is reliant upon their proximity, and facilitated by complex and elaborate astrocyte morphologies, we evaluated how loss of Slc22a3 impacts astrocyte morphology. Furthermore, Slc22a3-cKO astrocyte demonstrate downregulation of several morphology-related genes (**Fig 4E**). To assess astrocyte morphology, we performed high resolution confocal imaging coupled with IMARIS software analysis to enable three-dimensional reconstruction of OB astrocytes using GFP reporter in control (*Aldh1l1-GFP; Slc22a3-FF* + pAAV-Gfap-mCh) and Slc22a3-cKO (*Aldh1l1-GFP; Slc22a3-FF* + pAAV-Gfap-Cre-RFP) OB’s (**Fig. 4F**). Sholl analysis of astrocytes revealed decreased morphological complexity in Slc22a3-cKO in comparison to controls (**Fig. 4G**) based on a host of parameters including number of process intersections as a function of distance from (**Fig. 4H**), total process length, number of branches and terminal points (**Fig. 4I**).

**Figure 4.**
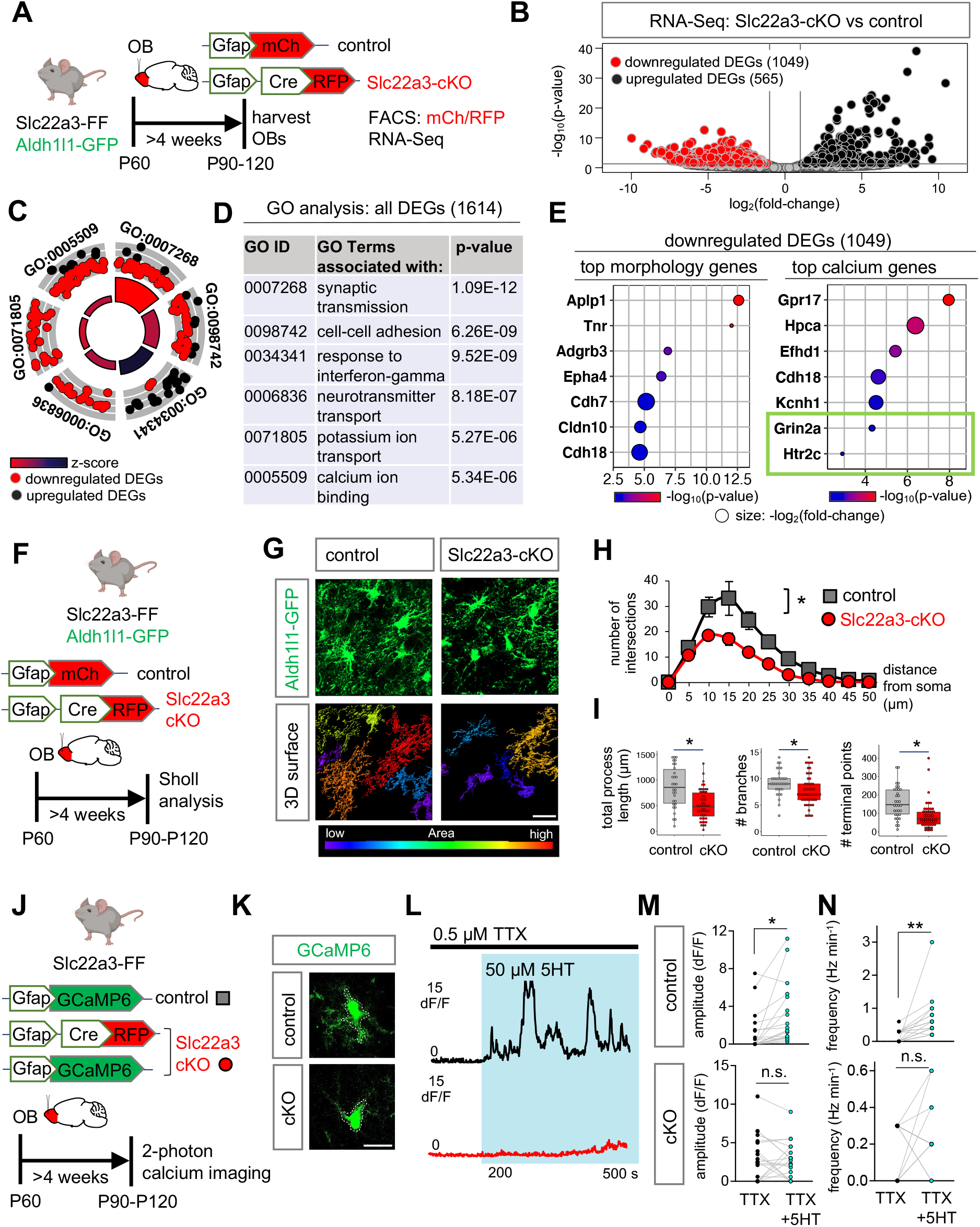
Astrocytic Slc22a3 regulates astrocyte morphology and calcium activity. (**A**) Schematic illustrating viral vectors and timelines for RNA-Seq experiment. (**B**) Volcano plots depicting RNA-Seq from FACS sorted astrocytes comparing Slc22a3-cKO vs. control samples. (**C-D**) Gene Ontology circle plot and table, showing the top GO terms found in DEGs shown in (B). (**E**) Top morphology and calcium associated genes (*p*< 0.01) in downregulated DEGs in the Slc22a3-cKO OB astrocytes. (**F**) Schematic illustrating viral vectors and timelines for evaluation of astrocyte morphology. (**G**) High-magnification confocal images of Aldh1l1-GFP from control and Slc22a3-cKO mice and 3D surface rendering of the same showing reduced astrocyte morphological complexity in Slc22a3-cKO OB astrocytes. Scale bar: 20 μm. (**H**) Sholl analysis of astrocyte complexity (n= 4, average of 34-54 cells/cohort, *p= 0.0117; two-way repeated measures ANOVA with Sidak correction). Data presented as mean ± SEM. (**I**) Quantification of total process length (\**p*= 0.0285, Wilcoxon rank sum test), branch number (\**p*= 0.0198, unpaired Student’s two-tailed t-test), and terminal points (\**p*= 0.0177, unpaired Student’s two-tailed t-test) in control and Slc22a3-cKO OB astrocytes (34-54 cells/cohort, statistics on n= 4 mice/cohort). (**J**) Schematic illustrating mice, viral vectors, and timelines for expression of optical calcium sensor in OB astrocytes. (**K-L**) Representative traces from two photon, slice imaging of GCaMP6 activity from OB astrocyte soma in control and Slc22a3-cKO in the presence of TTX (0.5 μM) and serotonin (5HT, 50 μM). Scale bar: 10 μm. (**M-N**) Quantification of amplitude and frequency from 5HT induced calcium activity from astrocyte soma (19-20 cells/cohort, control amplitude \**p*= 0.0346; Slc22a3-cKO amplitude *p*= 0.6992; control frequency \*\**p*= 0.0029; Slc22a3-cKO frequency *p*= 0.5588; paired Student’s two-tailed t-test on n= 4 mice/cohort).

Next, we evaluated astrocyte Ca^2+^ activity, as it is considered a proxy for the physiological activities of astrocytes and has been linked to astrocyte regulation of neuronal function. Furthermore, our RNA-Seq studies identified several genes linked to Ca^2+^ activity demonstrating significant downregulation in Slc22a3-cKO astrocytes (**Fig. 4E**). To assess astrocytic calcium, we used pAAV containing Gfap promoter driven GCaMP6 encoding a fluorescent calcium optical sensor, which we injected into OBs of control (Slc22a3-FF) and Slc22a3-cKO (Slc22a3-FF + pAAV-Gfap-Cre) mice (**Fig. 4J**). We generated OB slices from these mice and performed two-photon imaging to assess calcium dynamics under basal conditions from astrocytes located in the granular cell layer of the OB, finding no change in soma spontaneous calcium amplitude and frequency between control and Slc22a3-cKO OB astrocytes (**Supp Fig. S14A-C**).

Prior studies have shown that astrocyte Ca^2+^ activity is modified in response to neurotransmitters and neuromodulators providing an indirect measure of their interactions with neurons (*16*). Among the neurotransmitter and neuromodulator receptors that can influence Ca^2+^ activity, we observed a significant downregulation of the glutamate receptor *Grin2a* and serotonin receptor *Htr2c* in Slc22a3-cKO OB astrocytes (**Fig. 4E**). Therefore, to measure Ca^2+^ responses mediated by these receptors we assessed calcium activity in astrocytes from OB slices after application of glutamate and serotonin. Bath-applying serotonin or glutamate onto OB slices while monitoring Ca^2+^ elevations in astrocytes (with TTX in the solution) is a method to test astrocyte 5HT or Glu receptor (Gq or Gi)-mediated Ca^2+^ events, and associated astrocytic receptor activity. Furthermore, because Slc22a3 has been implicated in maintenance of extracellular serotonin levels in the brain (*38*), we were especially interested in serotonin induced Ca^2+^ dynamics. Similar to above, two-photon imaging was performed in control and Slc22a3-cKO mice, but TTX (0.5 μM) was applied to block neuronal activity for 200s before application of glutamate (300 μM) or serotonin (50 μM) for an additional 300s. With serotonin, quantification of fluorescence from astrocytic soma revealed a significant reduction in serotonin-induced amplitude and frequency of calcium events in Slc22a3-cKO OB astrocytes in comparison to controls (**Fig. 4K-N**). In contrast, with glutamate, we observed no significant difference between amplitude of glutamate-induced calcium activity in Slc22a3-cKO astrocytes in comparison to controls, whereas frequency showed a small reduction in Slc22a3-cKO (**Supp Fig. S14D-F**).

Analysis of astrocyte microdomain calcium activity in response to serotonin or glutamate revealed reduced amplitude and frequency of calcium events for both in Slc22a3-cKO (**Supp Fig. S15A-D**). Taken together, the observed reductions in astrocyte morphological complexity and calcium activity, indicate that astrocyte-neuron communication is impaired in astrocytic Slc22a3-deficient OBs, and this deficit is likely driven by impaired neuronal signaling.

### Slc22a3 regulates histone serotonylation in astrocytes

Because Slc22a3 is implicated in the transport of serotonin, we next examined intracellular serotonin levels within astrocytes in the OB. (**Supp Fig. S16A**). Immunostaining for serotonin (5-HT) revealed a decrease in its intensity within Slc22a3-cKO astrocytes (**Supp Fig. S16B-C**), suggesting a defect in serotonin transport. Recent studies in neurons have shown that serotonin can be added to histones and that serotonylation of histones directly potentiates epigenetic mechanisms of gene regulation (*31*). While both astrocyte morphology and calcium have been linked to a host of behavioral outcomes (*39*), the role of astrocytic histone modifications and associated epigenomic phenomenon in the regulation of circuits and behavior remains unknown. Furthermore, Slc22a3 is a serotonin transporter in the central nervous system and the role of serotonin in olfactory bulb astrocytes remains relatively unexplored, which led us to investigate whether serotonylation of histones regulates astrocyte function and associated olfactory circuits. We first asked whether the observed reduction in intracellular serotonin in Slc22a3-cKO astrocytes impacts histone serotonylation. Using the Aldh1l1-GFP reporter mouse, we performed immunostaining with antibodies specific for the serotonylated histone mark on histone H3, H3K4me3Q5ser (H3-5HT) and found that 86% of GFP+ OB astrocytes contain H3-5HT (**Fig. 5A**; **Supp Fig. S17A**). Next, we immunostained for H3-5HT in Slc22a3-cKO (*Aldh1l1-GFP; Slc22a3-FF* + pAAV-Gfap-Cre) and control (*Aldh1l1-GFP; Slc22a3-FF* + pAAV-Gfap-mCh) OB astrocytes (**Fig. 5B**), finding a reduction in the levels of H3-5HT in Slc22a3-cKO OB astrocytes (**Fig. 5C-D**), but not in neurons of Slc22a3-cKO OB’s (**Supp Fig. S17B**). We next performed ChIP-Seq of H3-5HT on control and Slc22a3-cKO OBs to determine whether patterns of H3-5HT modification are affected across the epigenome after astrocytic Slc22a3 loss and observed a 2.3-fold reduction in H3-5HT peaks in Slc22a3-cKO OBs (**Fig. 5E**). These findings corroborate our immunostaining results and support the notion that OB astrocytes from Slc22a3-cKO mice exhibit altered levels of serotonin.

**Figure 5.**
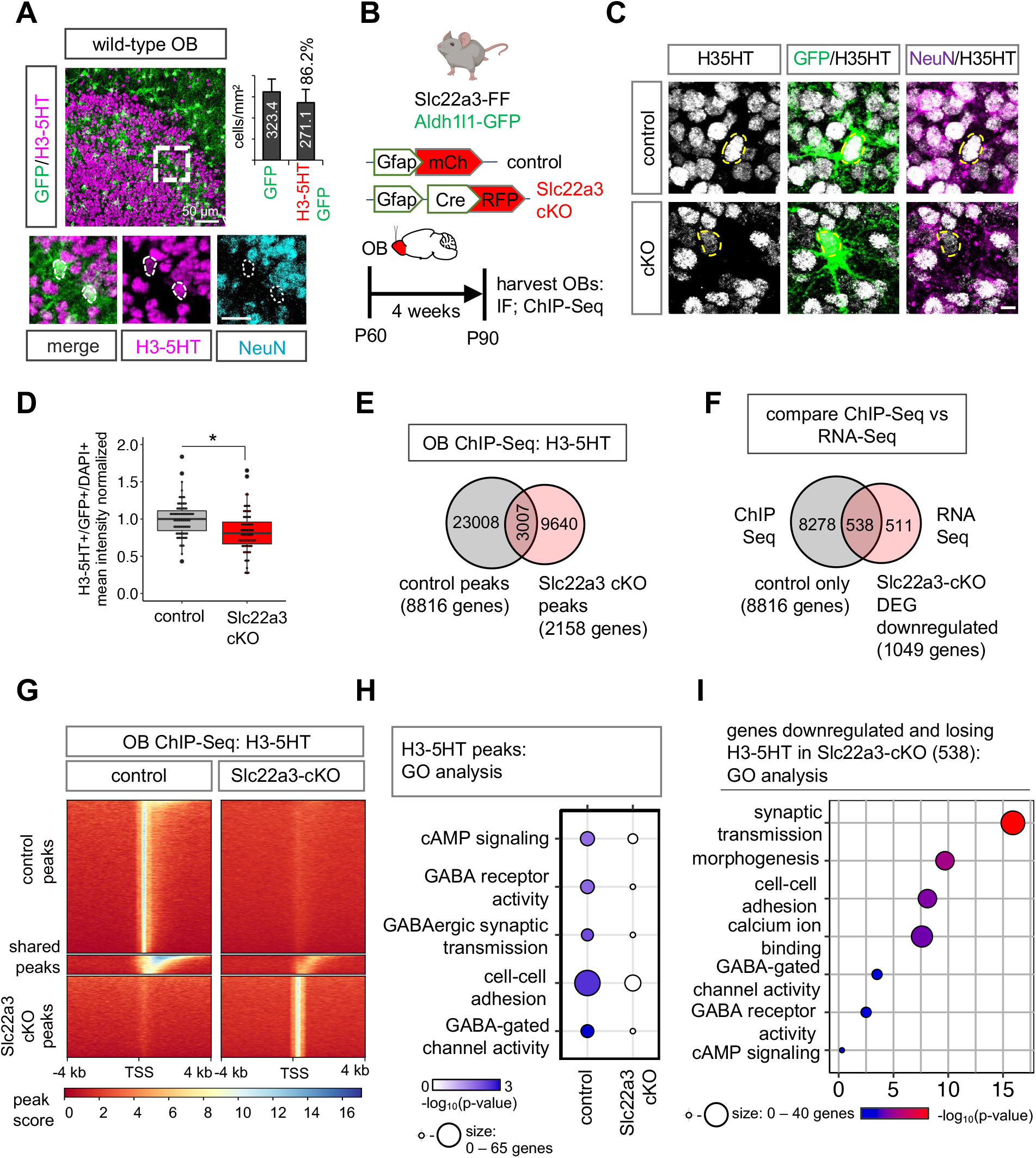
Slc22a3 regulates histone serotonylation in OB astrocytes. (**A**) Immunostaining of H3-5HT in OBs of Aldh1l1-GFP mouse and quantification of GFP+/H3-5HT+ co-labeling (n= 3, 25-45 cells). (**B**) Schematic illustrating viral vectors and timelines for H3-5HT quantification and ChIP-Seq. (**C-D**) H3-5HT immunostaining and quantification in control and Slc22a3-cKO OB astrocytes (74-79 cells/cohort, \**p*= 0.0377; unpaired Student’s two-tailed t-test on n= 4 mice/cohort). Scale bar: 5 μm. (**E**) Venn diagram depicting number of H3-5HT ChIP-Seq peaks unique and shared between control and Slc22a3-cKO OB’s (n=4 OBs/cohort). (**F**) Venn diagram depicting number of genes that both lose H3-5HT peaks and are downregulated in Slc22a3-cKO. (**G**) Heatmaps comparing ChIP H3-5HT at 4 kb from peak center in control vs Slc22a3-cKO OB’s. (**H**) GO analysis of genes at H3-5HT peaks revealing loss of H3-5HT regulation at GABA-associated pathways in Slc22a3-cKO, and (**I**) of the 538 overlapping genes shown in Figure 5F.

Prior studies demonstrating that histone serotonylation activates gene expression (*31*), led us to cross-compare our ChIP-Seq results with the RNA-Seq data obtained from Slc22a3-cKO, described earlier. Strikingly, this revealed that of the 1049 downregulated DEGs in the Slc22a3-cKO, 538 genes (51% of the DEGs) also lose histone serotonylation modifications in the Slc22a3-cKO (**Fig. 5F-G**). In comparison, of the 565 upregulated DEGs in Slc22a3-cKO, only 26 genes (4.6% of the DEGs) gain H3-5HT modifications in the Slc22a3-cKO (**Supp Fig. S17C**). We next interrogated gene ontologies and pathways exhibiting differential H3-5HT modification in the OB from Slc22a3-cKO mice. GO analysis between control and Slc22a3-KO H3-5HT peaks revealed that a prevalence of GO terms linked to GABAergic signaling are lost after astrocytic Slc22a3 deletion (**Fig. 5H**). Among these genes are various receptor subtypes for GABA, and GABA biosynthetic enzymes *Maob* and *Aldh1a1* (**Supp Fig. S17D**). Further analysis of the GO terms from the 538 genes that lose H3-5HT and are downregulated in Slc22a3-cKO revealed categories of synaptic transmission, morphology, cell-cell adhesion and calcium ion binding, GO terms that were also observed in the Slc22a3-cKO RNA-Seq (**Fig 4D**). In addition, GABA signaling pathway GO categories were also maintained in this smaller subset of filtered astrocytic genes (**Fig. 5I**).

Since the GABA pathway represents one of the top GO terms demonstrating a loss of histone serotonylation modifications in the Slc22a3-cKO (**Fig. 5H**), we therefore focused on the GABA synthesis pathway. Assessing astrocyte-specific expression of GABA and protein expression of *Maob* and *Aldh1a1* using localization with GFP in control and Slc22a3-cKO, we found that both MAOB and astrocytic GABA were significantly downregulated in the OB from Slc22a3-cKO mice, while Aldh1a1 was unaffected (**Fig. 6A-D**; **Supp Fig. S18A-B**). Similar analysis evaluating GABA and MAOB levels in neurons with NeuN colabeling revealed no significant differences (**Fig. 6E-H**). However, we observed overall reduction in mean intensity of GABA in whole sections, but not of MAOB (**Supp Fig S18C-D**). Given that astrocytes are capable of synthesizing and releasing GABA (*25, 40*) and our results suggest a general decrease in astrocytic GABA in Slc22a3-cKO astrocytes, we next used slice electrophysiology recordings to measure tonic GABA release from OB astrocytes in control and Slc22a3-cKO mice (**Fig. 6I**). Strikingly, we found a drastic decrease in tonic GABA current from granular cell layer neurons of the OB in Slc22a3-cKO. To exclude the possibility that the tonic GABA current is derived from neurons, we performed control recordings in the presence of TTX (0.5 μM) to dampen neuronal activity, which also revealed a decrease in tonic current in Slc22a3-cKO OBs, thus confirming that the observed reduction in tonic GABA current is driven by astrocytes, and not neurons (**Fig. 6J-L**). Finally, we recorded tonic GABA current in the presence of saturating concentrations of GABA (5 μM) and identified no differences between control and Slc22a3-cKO groups, further indicating that the observed reduction in tonic GABA is not driven by synaptic GABA receptors (**Fig. 6M-N)**. Collectively, these results indicate that Slc22a3 regulates histone serotonylation in OB astrocytes, which epigenetically controls astrocytic GABA and subsequent tonic GABA release by astrocytes.

**Figure 6.**
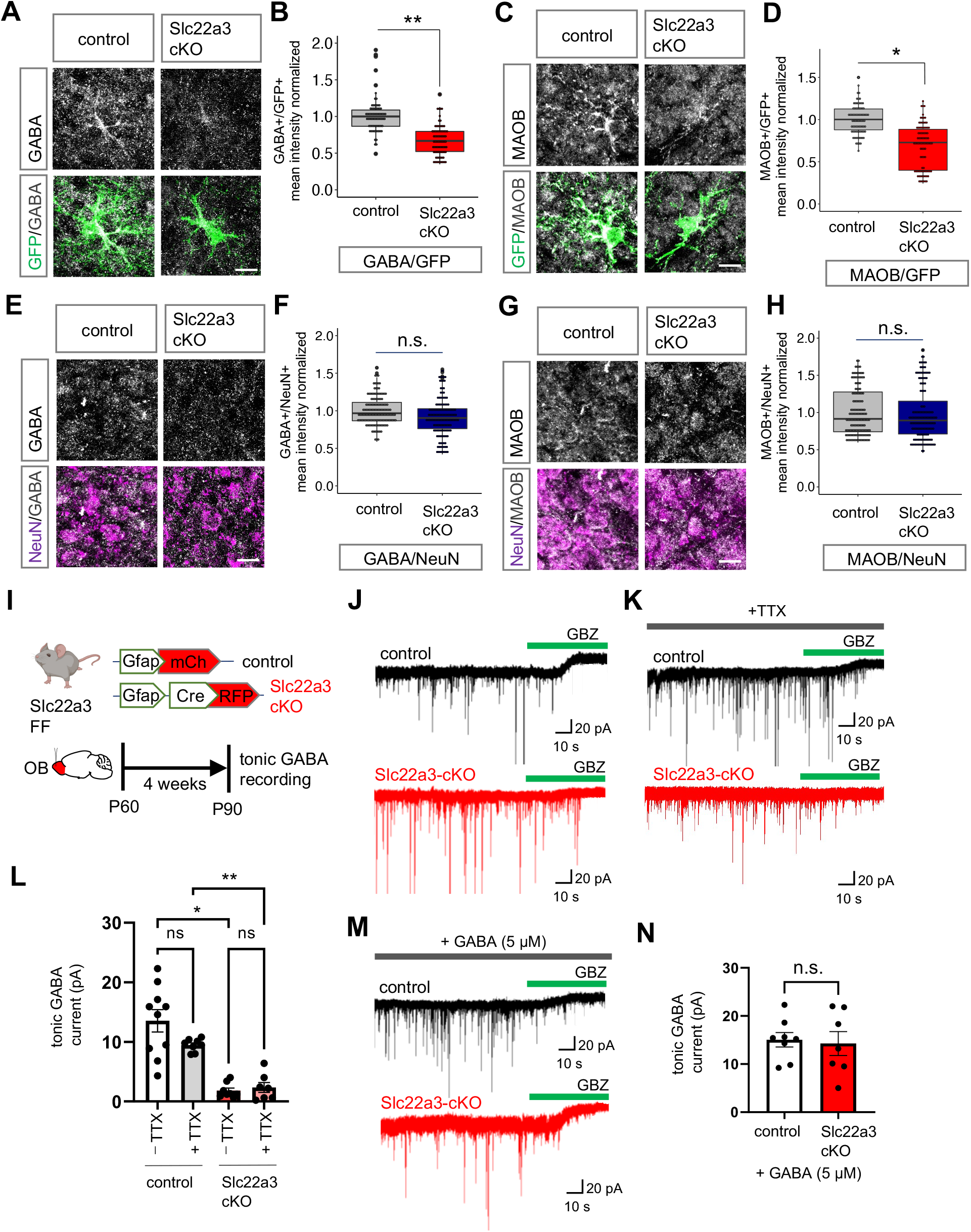
Slc22a3 regulates tonic GABA in OB astrocytes. (**A-H**) Immunostaining and quantification of GABA and MAOB in control vs. Slc22a3-cKO (A-D) GFP astrocytes (73-88 cells/cohort; GABA \*\**p*= 0.0032, MAOB \**p*= 0.0121) and (E-H) NeuN+ neurons in the OB (144 cells/cohort; GABA *p*= 0.2119, MAOB *p*= 0.9355); unpaired Student’s two-tailed t-test on n= 4 mice/cohort. Scale bar: 10 μm. (**I**) Schematic illustrating viral vectors and timelines for tonic GABA experiments. (**J-K**) Representative traces of tonic GABA currents in granule cells in OBs from control and Slc22a3-cKO treated with gabazine (GBZ, 20 μM), with or without TTX (0.5 μM). (**L**) Quantification of tonic GABA current (7-10 cells/cohort, –TTX \**p*= 0.0331, +TTX \*\**p*= 0.0056) (**M-N**) Representative traces and quantification of measurement of tonic GABA current in presence of GABA (7-8 cells/cohort, *p*= 0.8483); unpaired Student’s two-tailed t-test on n= 3 mice/cohort. Data presented as mean ± SEM.

### Inhibition of H3-5HT in OB astrocytes disrupts sensory processing

To examine whether H3-5HT modifications in OB astrocytes directly contribute to olfactory sensory processing, we utilized a recently published dominant negative mutant H3.3Q5A, where the histone glutamine residue that is modified by serotonin is mutated to an alanine to attenuate H3-5HT (*31*). The use of histone variant 3.3 (H3.3) ensures histone incorporation to replace canonical H3 during histone protein turnover (*41*). We generated pAAV containing control H3.3 and mutant H3.3Q5A under control of the Gfap promoter and introduced these into the OBs of wild-type mice (**Fig. 7A**). Astrocyte-specific expression of H3.3 and H3.3Q5A was confirmed by colocalization of the GFP reporter on H3.3 constructs with astrocyte marker Sox9 and neuronal marker NeuN (**Supp Fig. S19A-B**). We also observed no differences in cell numbers of GFP+ astrocytes and NeuN+ neurons in OB’s expressing these H3.3 constructs (**Supp Fig. S19C-D**). Quantification of H3-5HT revealed a significant reduction of H3-5HT in H3.3Q5A expressing OB astrocytes in comparison to H3.3 controls (**Fig. 7B-C**), but not in surrounding neurons (**Supp Fig. S19E**). Using this astrocytic H3-5HT attenuation mouse model, we next assessed astrocyte morphology and neuronal electrophysiology. As described earlier, high resolution confocal imaging, coupled with IMARIS software assisted 3D reconstruction of images and Sholl analysis revealed decreased morphological complexity in H3.3Q5A-expressing astrocytes compared to H3.3 controls (**Fig. 7D**) as evaluated by number of process intersections as a function of distance from soma (**Fig. 7E**), total process length, number of branches and terminal points (**Fig. 7F**). These data suggest that attenuation of astrocytic H3-5HT in the OB reduces astrocyte morphological complexity. Prior studies established that astrocytes exhibit extensive structural plasticity in perisynaptic astrocyte processes (PAPs) (*42, 43*), therefore we next measured the number of astrocyte terminal process points and determined distance between PAPs labeled by ezrin (Ezr) and postsynaptic marker PSD95. This analysis revealed a significant reduction in the number of PSD95 from Ezr+ terminal process points in H3.3Q5A-expressing astrocytes compared to control (**Supp Figure S20A-C**), suggesting reduced interactions between astrocyte processes and neuronal synapses. Together, these data suggest H3-5HT epigenetic processes regulate astrocyte morphology and astrocyte-neuron proximity in the OB.

**Figure 7.**
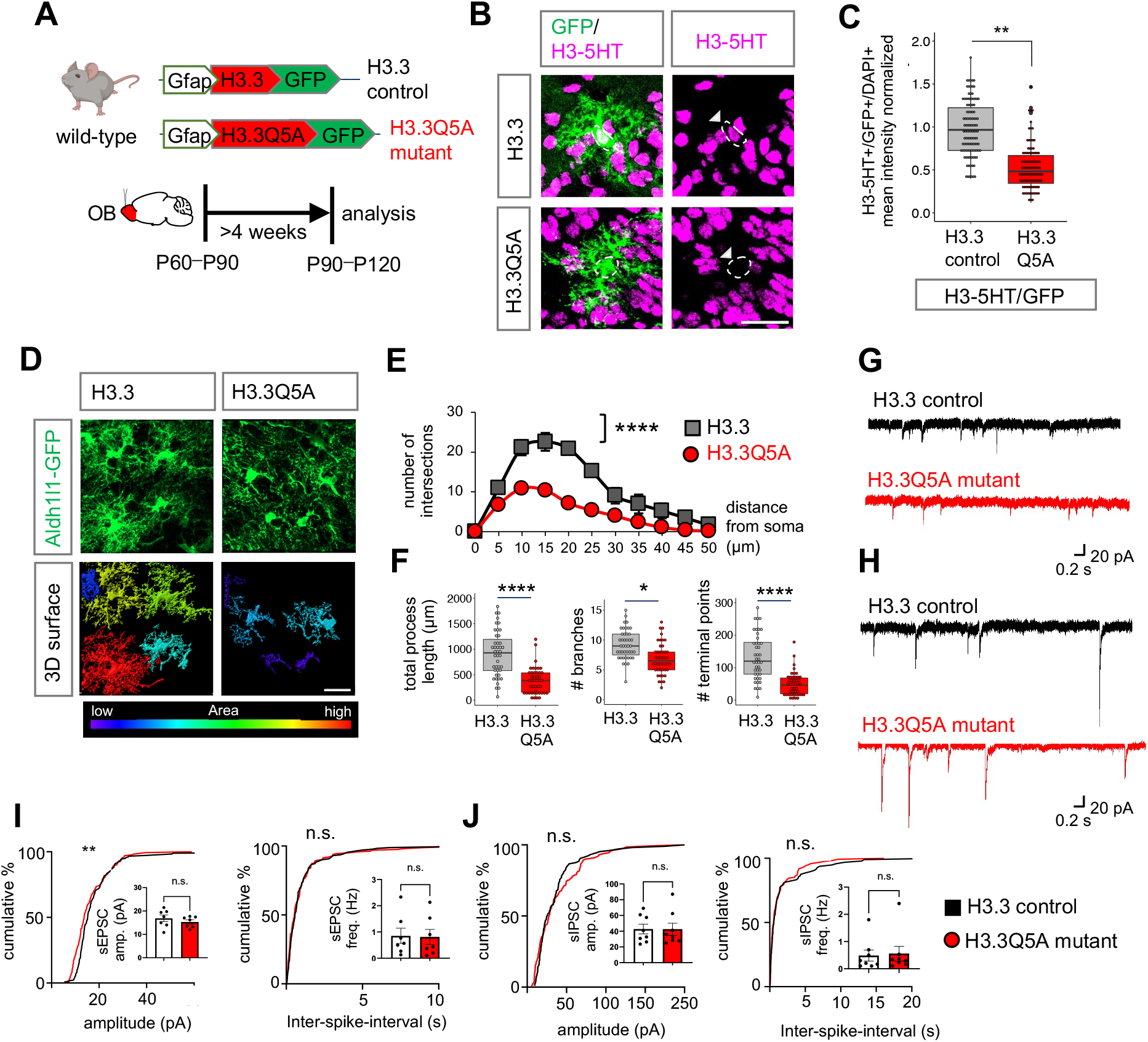
Inhibition of H3-5HT in OB astrocytes disrupts astrocyte morphology. **(A)** Schematic illustrating viral vectors used for H3.3 and H3.3Q5A expression in OB astrocytes. **(B)** H3-5HT and GFP co-labeling in H3.3 and H3.3Q5A expressing OBs and (**C**) box plots depicting quantification of astrocytic H3-5HT (99-110 cells/cohort, \*\**p*= 0.0092; unpaired Student’s two-tailed t-test on n= 4 mice/cohort). Scale bar: 25 μm. (**D**) High-magnification confocal images of H3.3-GFP and 3D surface rendering of the same showing reduced astrocyte morphological complexity in H3.3Q5A OB astrocytes. Scale bar: 20 μm. (**E**) Sholl analysis of astrocyte complexity (n= 4, average of 44 cells/cohort, ****p= 1.9e-05; two-way repeated measures ANOVA with Sidak correction). Data presented as mean ± SEM. (**F**) Quantification of total process length, branch number, and terminal points (44 cells/cohort, \*\*\*\**p*= 2.75e-05, \**p*= 0.0122, \*\*\*\**p*= 8.73e-06, unpaired Student’s two-tailed t-test on n= 4 mice/cohort). (**G-H**) Representative traces and (**I-J**) summary data of amplitude and frequency from (I) sEPSC recordings (7 cells/cohort, sEPSC amplitude *p=* 0.2386; sEPSC frequency *p*= 0.7917; unpaired two-tailed Student’s t-test on n= 3 mice/cohort, \*\**p*= 0.0059 K-S test); and from (J) sIPSC recordings (8 cells/cohort, sIPSC amplitude *p*= 0.8277, sIPSC frequency *p*= 0.7128, unpaired two-tailed Student’s t-test on n= 3 mice/cohort. All recordings are in granule cells from H3.3 and H3.3Q5A OB’s and data is presented as mean ± SEM.

Examination of basal properties of excitatory (**Fig. 7G**) and inhibitory (**Fig. 7H**) postsynaptic currents revealed no changes in sEPSC/sIPSC frequency and amplitudes (**Fig. 7I-J**). These data suggest that attenuation of astrocytic H3-5HT in the OB has no effect on spontaneous synaptic currents. Since Slc22a3-cKO showed a significant reduction in H3-5HT epigenomic regulation of GABA pathways, we next assessed expression of MAOB and GABA, finding a significant reduction of both in OB astrocytes expressing H3.3Q5A in comparison with H3.3 control (**Fig. 8A-C**), but not in neurons (**Supp Fig. S21A-B**). We also observed an overall reduction in GABA mean intensity in whole fields (**Supp Fig. S21C-D**). Furthermore, we performed additional control experiments to confirm that expression of Gfap-H3.3 alone does not alter levels of Slc22a3, GABA and MAOB, in comparison to Aldh1l1-GFP animals (**Supp. Fig. S21E-G**). To determine whether H3-5HT regulation of GABA production culminates in altered GABA release from OB astrocytes, we used electrophysiology to measure GABA release from OB astrocytes expressing H3.3Q5A mutant or H3.3 controls, finding a striking reduction in tonic GABA current in H3.3Q5A condition (**Fig. 8D-F**). Control recordings were performed in the presence of TTX (0.5 μM), revealing a decrease in tonic current in H3.3Q5A-expressing OBs consistent with alterations astrocyte-derived tonic GABA currents. Additional control recordings in the presence of saturating GABA (5 μM) further corroborate reduced tonic GABA currents in OB astrocytes expressing H3.3Q5A (**Supp. Fig. S22A-B**).

**Figure 8.**
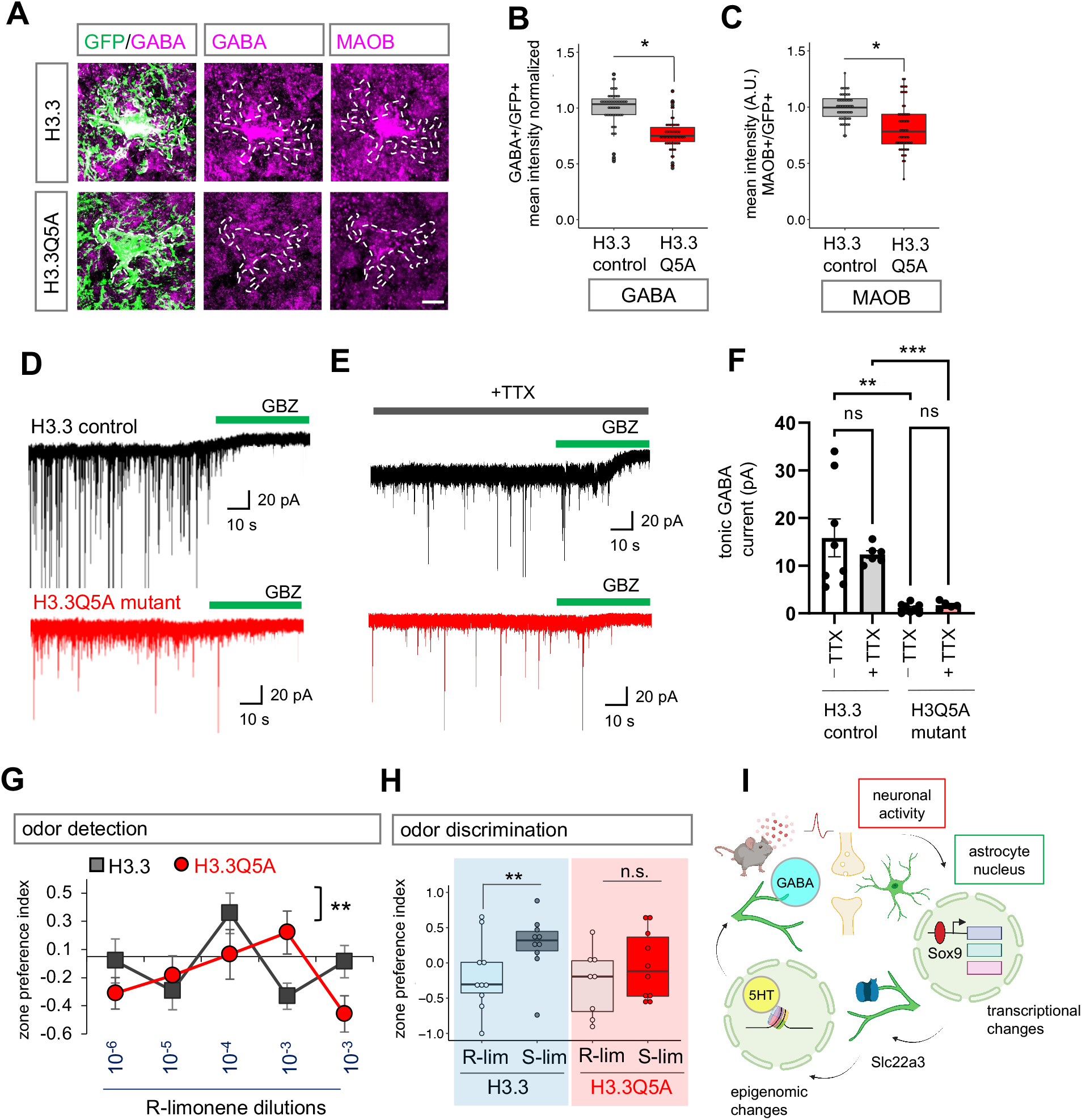
Inhibition of H3-5HT disrupts astrocytic tonic GABA and sensory processing. (**A-C**) Immunostaining and box plots depicting quantification of astrocytic GABA and MAOB in H3.3 and H3.3Q5A OBs (56-58 cells/cohort, GABA: \**p*= 0.0117, unpaired two-tailed Student’s t-test on n= 4 mice/cohort; MAOB \**p=* 0.0285, Wilcoxon rank sum test on n= 4 mice/cohort). Scale bar: 10 μm. (**D-E**) Representative traces of tonic GABA currents in granule cells in OBs from H3.3 and H3.3Q5A treated with gabazine (GBZ, 20 μM), with or without TTX (0.5 μM). (**F**) Quantification of tonic GABA (5-8 cells/cohort, –TTX \*\**p*= 0.0017, +TTX \*\*\**p*= 0.0007; unpaired two-tailed Student’s t-test on n= 3 mice/cohort). Data presented as mean ± SEM. (**G**) Quantification of odor detection in H3.3 and H3.3Q5A mice (n=10/cohort, \*\**p*= 0.0072; two-way repeated measures ANOVA with Sidak multiple comparison). (**H**) Quantification of odor discrimination between R-lim and S-lim from the same cohorts of mice (n=10/cohort, **p= 0.0058; two-way repeated measures ANOVA with Sidak multiple comparisons). (**I**) Model figure integrating activity dependent transcriptional changes in astrocytes, with Slc22a3 function in olfactory circuits, and histone serotonylation regulation of GABA in astrocytes.

Given the links between GABA signaling and neuronal circuit function in the OB, we next examined how inhibition of H3-5HT impacts olfactory sensory processing. Towards this, we performed odor detection and odor discrimination behavioral assays as previously described (**Fig. 3**). First, we compared odor behaviors of mice expressing H3.3 in OB astrocytes with wild-type control mice to assess whether astrocytic H3.3 expression alone effects odor behavior, finding that it has no significant impact on odor detection limit and odor discrimination ability (**Supp Fig. S22C-D**). Next, we compared odor behaviors of H3.3 control-and H3.3Q5A mutant-expressing mice, which revealed reduced odor detection and impaired odor discrimination in mice expressing H3.3Q5A in OB astrocytes (**Fig. 8G-H**). Strikingly, mice expressing astrocytic H3.3Q5A recapitulated the phenotypes of impaired tonic GABA and olfactory behaviors observed in the Slc22a3-cKO (**Figs. 3-4, 6**). Collectively, these findings suggest that the neuronal activity-dependent target Slc22a3 in OB astrocytes facilitates serotonin mediated epigenetic regulation of GABA synthesis, which contributes to sensory processing of olfaction (**Fig. 8I**).

## Discussion

Neuronal circuits and associated behaviors are intimately linked to astrocytes, yet how neuronal activation sculpts astrocyte transcriptional responses to support circuit activities is unclear. Using olfactory sensory processing as a model for neuronal activation, we identified neuronal activity-dependent transcriptional changes in astrocytes, highlighted by alterations in DNA-binding capacity of the transcription factor Sox9 and induction of a prospective immediate early gene, the neuromodulator transporter Slc22a3. Further examination of Slc22a3 in astrocytes revealed that it is required to maintain olfactory circuit function. Our study pinpointed Sox-family TFs as key components of activity-dependent responses in astrocytes in the OB. However, given that astrocyte function is regulated by region-specific transcriptional dependencies (*13, 14, 44*), there are likely to be region-specific astrocyte TFs that mediate these responses and responses associated wtih injury and disease (*45*). Furthermore, it is also likely that different forms of neuronal input (excitatory, inhibitory, etc.) or behavioral states influence these transcriptional responses in astrocytes. These findings illustrate that astrocytes exhibit a form of plasticity in response to neuronal activity and this plasticity is reliant upon a combination of inputs from neuronal circuits and region-specific TFs. These observations also raise the question of how this form of transcriptional plasticity in astrocytes is regulated. The increase in Sox9-DNA binding in the presence of heightened neuronal activity is likely the result of activity-dependent protein interactions. Our Mass-Spec studies identified a cohort of 40 proteins that interact with Sox9 in the OB specifically after neuronal activation. These findings indicate that neuronal input influences the protein constituency of transcriptional complexes and suggests that these inputs shape protein interactions that drive gene expression. Understanding how neuronal input remodels transcriptional complexes and the specific roles of activity-dependent protein interactions in astrocyte-neuron communication are important areas of future investigations. From these collective findings a model emerges, where neuronal activity orchestrates transcriptional responses in astrocytes to meet the demands of a functioning circuit.

We identified Slc22a3 as a prospective immediate early gene that is specifically induced in OB astrocytes after exposure to odor, and analysis of OB astrocytes from Slc22a3-cKO mouse revealed a host of phenotypes including reduced morphological complexity, reduced Ca^2+^ activity, decreased tonic GABA release, altered histone serotonlyation, and impaired olfactory detection. These observations suggest that in addition to an acute function after odor exposure or neuronal activity, Slc22a3 also plays a chronic, longer-term role in astrocyte function and communication with neurons. The chronic role is likely due to epigenomic changes that result in increased MAOB expression, GABA production, and altered morphology. This chronic role is likely to reflect a homeostatic function for Slc22a3 and/or serotonin transport in maintaining physiological activities of OB astrocytes. It is also possible that these two roles are interdependent, where the changes in GABA release from astrocytes influences neuronal activity in a way that impacts astrocyte morphology. Alternatively, we must also consider the possibility that shorter astrocytic processes may lead to longer diffusion time for GABA to reach synaptic GABA receptors, ultimately leading to reduced tonic GABA current. Another possibility is that there is overall less tonic GABA release from astrocytes, as suggested from our data of reduced astrocytic GABA biosynthetic enzyme MAOB and astrocytic GABA levels. Slc22a3 functions as a monamine transporter that has been shown to regulate the transport of serotonin (*38*). Because the OB is densely innervated by serotonergic fibers, which activate interneuron granule cells (*46*), it raises the question of whether serotonin is a factor that modulates inhibitory outputs and the subsequent impact on odor discrimination. Our finding that odor-evoked neuronal activation leads to enrichment of cAMP signaling GO terms in astrocytes, suggests that neuronal activation leads to astrocytic activation of Gs or Gi GPCRs, including astrocytic serotonergic receptors. In turn, this leads to upregulated expression of serotonin transporter Slc22a3 in astrocytes via Sox9. Our data demonstrating serotonin-induced calcium signaling is reduced in Slc22a3-deficient astrocytes is further supported by our observation of reduced astrocytic serotonergic receptors in Slc22a3-cKO. These collective findings indicate that astrocytes utilize serotonin through Slc22a3 to control tonic GABA inhibition, highlighting a new mechanism of serotonergic modulation of inhibitory outputs through astrocytes, which ultimately impacts olfactory sensory processing.

Epigenetic mechanisms of gene regulation play a key role in all facets of cell physiology, but how these processes influence roles for astrocytes in circuits remains undefined. We found that serotonin is added to histones in astrocytes and this modification is utilized to regulate gene expression and olfactory sensory processing. Mechanistically, Slc22a3 regulates serotonin levels in astrocytes, which influences the extent of histone serotonylation, reflecting a long-term change in transcriptional activity that ultimately regulates the expression of GABA-synthesis components. This represents a new mechanism of epigenomic regulation in astrocytes, while also highlighting that a host of epigenomic phenomenon remain undefined in astrocytes. A recent study demonstrated that Slc22a3 may have roles in noradrenergic signaling at astrocyte nuclear membranes, highlighting another way for Slc22a3 to modulate nuclear processes (*47*). Synaptic changes in neurons affect chromatin accessibility (*30, 48, 49*), and it is likely that classic histone acetylation and methylation states are also modified in astrocytes by neuronal activity. Indeed, Sox9 has recently been implicated in regulating epigenomic states in brain tumors (*50*). In the context of olfactory circuits, a few studies have investigated the role of serotonergic system in modulation of olfactory processing (*51, 52*). Here, our studies reveal a new role for serotonin in astrocytes, wherein it gets deposited in the genome and indirectly regulates astrocytic release of GABA, demonstrating how astrocytes use serotonin to gate GABA in the olfactory bulb. A host of prior studies in mouse and other species have established links between neuromodulator (volume-based) and neurotransmitter (synaptic) signaling (*28, 29, 53, 54*), mediated by astrocytes. Our studies are distinct in that we identified an epigenomic intermediatory. It will be critical to decipher the extent to which these neuromodulatory signaling mechanisms in astrocytes directly regulate neurotransmitter signaling or go through an epigenomic intermediatory and the nature of this form of transcriptional regulation under different behavioral states.

## ACKNOWLEDGMENTS

We thank Anna Yu-Szu Huang for providing training in stereotaxis injection and Kevin Ung for comments regarding odor behavioral assays. Images in schematics were created using Biorender.com.

## Author Contributions

Conceptualization: D.S., B.D.; Methodology and data acquisition: Y.-T.C., J.W.; D.-J.C, Z.-F.L., W. K., H.-C.C., B.L., A.C., K.R., A.J., Resources: T.-W.H., B.A., I.M., and B.D.; Funding acquisition: D.S., B.D.; Writing original draft: D.S, B.D.; Review and editing: D.S., I.M., B.D.

## Data and materials availability

All data are available in the manuscript or supporting materials. The RNA-Seq and ChIP-Seq datasets generated in this study will be made available at NCBI GEO.

## SUPPLEMENTARY MATERIALS

Materials and Methods

Supporting Figures S1−S22

References (*55–68)*

