## Supporting Materials for "Activity-dependent induction of astrocytic Slc22a3 regulates sensory processing through histone serotonylation"

**This PDF file includes:**

Materials and Methods

Supporting Figures S1–S22

Supporting References (*55–68*)

### Materials and Methods:

Animals

All animals were treated in compliance with the US Department of Health and Human Services, NIH guidelines and Baylor College of Medicine IACUC guidelines. Mice were housed in a 12-hour light/dark cycle environment with food and water available at all times. Both male and female mice were used for all experiments and littermates were randomly allocated to experimental groups. All mice used in this study were maintained on the C57BL/6J background. For all *ex vivo* and *in vivo* experiments, adult mice aged 2-month to 5-month were used. For Sox9 conditional knockout, Sox9 flox/flox (FF) conditional mutant mice was crossed with Cag-CreER mice. Conditional deletion of astrocytic Sox9 has been characterized previously (*55*) and we followed the same protocol for Sox9 deletion (Supp Fig. S8A). For Slc22a3 conditional knockout, Slc22a3-FF conditional mutant mice was generated at the Baylor College of Medicine Genetically Engineered Mouse core facility, with floxed sites flanking exon 2. For FACS purification and visualization of astrocytes the Aldh1l1-GFP reporter mouse line was used (*56*). Sox9-FF and Slc22a3-FF was crossed with Aldh1l1-GFP for histological analysis in conditional knockout mice. All above experimental procedures were approved by Baylor College of Medicine IACUC.

Plasmids used for pAAV vectors

For chemogenetic experiments with Gq-DREADD, pAAV-hSyn-hM3D(Gq)-mCherry (Addgene plasmid #50474) was used. For Sox9 and Slc22a3 conditional knockout with Gfap-Cre-RFP, AAV-GFAP-iCre-P2A-TurboRFP (kind gift from Kevin Ung and Benjamin Arenkiel) was used (*55*). For controls with Gfap-mCh, pZac2.1-GfaAB1CD-mCherry was used. For two-photon calcium imaging experiments, we generated pAAV-GFAP-GCaMP6-GFP from flexed GCaMP6m and pZac2.1-GfaBC1D-mCherry-hPMCA2w/b (Addgene #111568). For experiments with H3.3 control and H3.3Q5A mutant, we generated pAAV-Gfap-H3.3/H3.3Q5A-GFP from H3.3(wild type or Q5A)-Flag-HA(C-Ter) (*57*) in a pAAV-IRES-GFP backbone, wherein CMV promoter was replaced with Gfap promoter from AAV-GFAP-iCre-P2A-TurboRFP using MluI and BamHI restriction sites. Purified plasmids were validated using Genewiz sequencing services. All viral expression vectors were packaged into AAV at the Viral Vector core facility at the Neurological Research Institute at Baylor College of Medicine.

Delivery of pAAV vectors into mouse brain by intraventricular or stereotaxis injection

For chemogenetic experiments, pAAV for Gq-DREADD (serotype 2/9, ~7.7E+12 genome copies/ml) was introduced by intraventricular injection into postnatal day 1-2 pups. Trypan Blue dye (2.5 μl) was mixed with AAV (10 μl) before injection. Pups were anesthetized with hypothermia and AAV mixed with dye (2 μl) was injected into each hemisphere. Pups were allowed to recover in cages placed partially on low-voltage heating pad for atleast 2 hours. Subsequent Gq-DREADD experiments was performed when pups reached adult age (>8 weeks). For region-specific AAV delivery into the OB, stereotaxis microinjection was performed in 2-3 month old mice. The following viral titers were used: Gfap-mCh (~3E+12 GC/ml), Gfap-Cre (~2.6E+12 GC/ml), Gfap-GCaMP6-GFP (~6.8E+12 GC/ml), Gfap-H3.3-GFP (~5.2E+12 GC/ml) and Gfap-H3.3Q5A-GFP (~3.71E+12 GC/ml); GC: genome copies. Fast Green dye (1 μl) was mixed with AAV (10 μl) before injection. Mice were injected intraperitoneally with Buprenex (0.3 mg/ml), anesthetized with isoflurane and placed into stereotaxic frame with the head secured by blunt ear bars and nose placed into anesthesia and ventilation system. Skin incision was made, followed by craniotomies of 2-3 mm in diameter above the OB powered by a high-speed drill. AAV was loaded into a micro dispenser (Nanoject II, Drummond Scientific, #13-681-460) and injected using beveled glass pipettes (Drummond Scientific, #3-000-203-G/X) at a rate of 69 nL/s, 10 s intervals, 10 times, in each olfactory bulb using the following co-ordinates:

4.7 mm anterior to bregma, 0.6 mm lateral to midline, depth of 1.5 mm beneath the surface of the skull. Glass pipettes were left in place for at least 3 min before and after AAV delivery prior to slow withdrawal. Surgical wounds were closed with sutures and mice were allowed to recover in cages placed partially on low-voltage heating pad for atleast 2 hours. Buprenex (0.3 mg/ml) was administered and mice were monitored for 3 days after surgery. All subsequent experimental analyses on these mice were performed 4-5 weeks after viral delivery. All animal procedures were performed in accordance with approved Baylor College of Medicine IACUC protocol.

In vivo neuronal activation

For chemogenetic neuronal activation (schematic shown in Fig. 1C), 2-3 month old Gq-DREADD expressing mice were administered clozapine-N-oxide CNO (Tocris #4936) at 0.3 mg/kg body weight intraperitoneally. After 30 minutes, mice were sacrificed and brains were collected. Control animals were injected with saline. Additional controls were injected with CNO at the same dosage in mice expressing empty viral vector. Only Gq-DREADD expressing mice that were administered CNO showed seizure-like behaviors within 10-15 minutes. For odor evoked neuronal activation, experimental paradigm was based on a previous report (*58*) and modified here as shown in schematic in Fig. 1J and Fig. 2I. Odorant isoamyl acetate was used since it is a standard non-biological and non-preferred odor. Control animals were exposed to mineral oil. Additional controls consisted of mice that were directly taken from home cages without habituation. For FACS purification and histological analyses, brains were collected as described below. For proteomic experiments, OBs were collected and snap frozen. For ChIP experiments, OBs were collected and processed immediately as described below.

Immunofluorescence on frozen brain tissues

Mice were anesthetized under isoflurane and perfused transcardially with PBS pH 7.4, followed by 4% paraformaldehyde (PFA), brains were removed, fixed in 4% PFA overnight, and placed in 20% sucrose overnight, and embedded in OCT the next day. Cryosections of 20 μm were washed in PBS twice, antigen retrieval was performed by incubating sections in 10 mM sodium citrate (pH 6.0; 0.05% Tween20) at 75°C for 10 min, blocked with 10% goat or donkey serum in PBS with 0.3% Triton X-100, and incubated with primary antibody dilutions in blocking solution overnight at 4°C. The next day, sections were washed in PBS with 0.1% Triton X-100 and incubated with species-specific secondary antibody dilutions for 1 hour at room temperature. After Hoechst nuclear counterstaining (1:10,000; ThermoFisher #H3570), coverslips were mounted with VECTASHIELD antifade mounting medium.

The following primary antibodies were used: mouse anti-mCherry (1:500, Abcam #ab125096), chicken anti-GFP (1:1000, Abcam #ab13970), rabbit anti-Cre (1:500, Cell Signaling #15036), rabbit anti-Slc22a3 (1:250, Alomone Labs #ACT-013), rabbit anti-Slc22a3 (1:100, Alpha Diagnostics #OCT-31A), rabbit anti-Fos (1:200, Cell Signaling #2250), rabbit anti-H3K4me3Q5ser (1:500. EMD Millipore #ABE2580), mouse anti-NeuN (1:500, EMD Millipore #MAB377), guinea pig anti-GABA (1:200, EMD Millipore #AB175), rabbit anti-MAOB (1:100, Proteintech #12602-1-AP), rabbit anti-Aldh1a1 (1:200, Abcam #ab52492), goat anti-serotonin (1:200, Abcam #ab66047), mouse anti-Gfap (1:500, EMD Millipore #MAB360), mouse anti-Ezrin (1:100, BioLegend #866401), rabbit anti-PSD95 (1:500, Invitrogen #51-6900).

The following secondary antibodies were used (1:500, 0.1% Triton X-100 in PBS): goat anti-mouse Alexa Fluor 568 (ThermoFisher #A11004), goat anti-mouse Alexa Fluor 488 (ThermoFisher #A11001), goat anti-mouse Alexa Fluor 647 (ThermoFisher #A21235), goat anti-chicken Alexa Fluor 488 (ThermoFisher #A11039), goat anti-rabbit Alexa Fluor 568 (ThermoFisher #A11036), goat anti-rabbit Alexa Fluor 488 (ThermoFisher #A11034), goat anti-rabbit Alexa Fluor 647 (ThermoFisher #A21244), goat anti-guinea pig Alexa Fluor 568 (ThermoFisher #A11075), donkey anti-goat Alexa Fluor 568 (ThermoFisher #A11057), donkey anti-chicken Alexa Fluor 488 (ThermoFisher #78948), donkey anti-mouse Alexa Fluor 647 (ThermoFisher #A31571).

Confocal imaging and analyses

Fluorescent images were acquired using a Zeiss LSM 980 confocal microscope with 20× or 40× oil objective. All images were taken from the granular cell layer of the OB in the region near internal plexiform layer (Supp Fig. S11B). We focused on this region since both Fos and Slc22a3 showed a robust expression pattern in this region. For quantification, both control and experimental groups were immunostained on the same day, images were acquired in one session on the same day and identical laser power settings were maintained for all cohorts under comparison. All quantification was done on raw unprocessed images. Quantification was done using ImageJ, wherein region of interest (ROI) was detected manually based on Aldh1l1-GFP labeled astrocyte boundary, and area or mean intensity values were extracted. Astrocytes for quantification were selected based on clear GFP reporter labeling and all analyses were done blind to conditions. Quantification of cell numbers was done using Cell Counter or Analyze Particles function in ImageJ. For visualization, box plots were made in R-studio using ggplot2.

FACS purification

Different mouse brain regions (cortex, hippocampus, olfactory bulb) were collected and dissociated using a previously described protocol (*59*). Dissociated cells were sorted on a BD FACSAria III instrument (100 μm nozzle). Around 95,000 cells were collected per 1.5 mL tube, which contained Buffer RLT (650 μl, Qiagen #79216) with 1% β-mercaptoethanol. Finally, each sample was vortexed and rapidly frozen on dry ice.

RNA extraction, library preparation and RNA-sequencing

Total RNA was extracted using the RNeasy Micro Kit (Qiagen #74004) and quality control was performed using the High Sensitivity RNA Analysis Kit (Agilent #472-0500) on a 12-capillary Fragment Analyzer. cDNA synthesis and library construction with 8-bp single indices were done from 10 ng total RNA using Trio RNASeq System (NuGEN #0507-96). The resulting libraries were validated using the Standard Sensitivity NGS Fragment Analysis Kit (Agilent #DNF-473-0500) and quantified using Quant-it dsDNA Assay kit (ThermoFisher #Q33120). Samples were diluted to equimolar concentrations (2 nM), pooled and denatured according to manufacturer's instructions. A final library dilution of 1.3 pM was subjected to paired-end (read 1: 75, read 2: 75) sequencing of approximately 40 million reads per sample using the High Output v2 kit (Illumina #FC-404-2002) on a NextSeq550 instrument.

RNA-sequencing bioinformatics analysis

Sequencing files from each flow cell lane were downloaded in fastq files and merged. Quality control was performed using fastQC (v0.10.1) and MultiQC (v0.9). Reads were mapped to the mouse genome mm10 assembly using STAR (v2.5.0a) (48). In R (v4.1.2), mapped reads were used to build count matrices using Bioconductor packages GenomicAlignments (v1.26.0) and GenomicFeatures. (v1.42.2) (*60*). University of California Santa Cruz transcripts were downloaded from Illumina iGenomes in the GTF file format. DESeq2 (v1.20.0) (*61*) was used for normalization and differential gene expression analysis. Motif analysis was performed using Hypergeometric Optimization of Motif Enrichment (HOMER, v4.10) to identify transcription factor motifs enriched within 1000 bp before and 500 bp after transcription start site. Motifs with $p< 0.05$ were selected and further filtered based on two criteria: (1) >1.5 fold expression in GFP astrocytes over mCh neurons; (2) transcript expression >150 CPM in GFP astrocytes. GOs were determined using Enrichr, and significant GO terms ($p< 0.01$) were selected. For visualization, plots were made using ggplot2 (v3.3.5). Gene expression heatmaps were generated using ComplexHeatmap (v2.6.2).

RT-qPCR

RNA extraction and cDNA preparation was performed as described above for sample preparation of RNA-Seq. RT-qPCR was performed using PerfeCTa SYBR Green Fast Mix (Quantabio #95072-012) on a Roche Light Cycler 480 instrument. Reactions were set up using 2 ng cDNA, 250 nM primers, and 1× SYBR mix. qPCR was carried out at 95°C for 30 s, 40 cycles of 95°C for 5 s and 60°C for 30 s, with subsequent melting curve analysis. The expression of transcripts of target genes was normalized to *Gapdh*. RT-qPCR primers used are as follows: for *Slc22a3* (forward 5'- GGAGACCCACTCTACCATCGT-3', reverse 5'-GCTGCATAGCCCAAGGTAAAA-3'); for *Gapdh* (forward 5'- TGGCCTTCCGTGTTCCTAC-3', reverse 5'- GAGTTGCTGTTGAAGTCGCA-3').

Chromatin immunoprecipitation (ChIP)

For ChIP-Sox9 and ChIP-Sox2 in the brain, we pooled cortices, hippocampi, and olfactory bulbs from 3 mice for each experimental cohort. For ChIP-Sox9 from odor exposed mice, we pooled olfactory bulbs from 6-8 mice for each experimental cohort. For ChIP-H35HT, we pooled olfactory bulbs from 4 mice for each experimental cohort and performed two independent sequencing runs. Immediately after harvesting, tissues were dissociated in cold PBS using a pellet homogenizer on ice. Chromatin was crosslinked using a freshly prepared 1.1% formaldehyde solution with rocking at room temperature for 10 min, followed by addition of 0.1 M glycine. Cell pellets were collected by centrifugation at 3,500 rpm for 5 min at 4°C, washed with PBS, and frozen at 80°C until further processing. Pellets were resuspended with PBS/PMSF containing 0.5% Igepal to release nuclei followed by washing with cold ChIP-Buffer (0.25% Triton-X100, 10 mM EDTA, 0.5 mM EGTA, 10 mM HEPES pH 6.5), and nuclei were lysed with ChIP lysis buffer (0.5% SDS, 5 mM EDTA, 25 mM Tris-HCl pH 8) for 15 to 20 min at room temperature. Lysates were sonicated to 250 to 350 bp using Diagenode Bioruptor. Chromatin was quantified using the Quant-iT double-stranded DNA (dsDNA) Assay kit (Thermo Fisher, #Q33120), diluted 5-fold (2 mM EDTA, 150 mM NaCl, 1% Triton X-100, 20 mM Tris-HCl; pH 8.0; with protease inhibitors), and incubated with antibody overnight at 4°C with rotation. For ChIP-Sox9, 70-100 µg of chromatin was incubated with rabbit anti-Sox9 (7-10 µg; Abcam #ab5535). For ChIP-Sox2, 70-100 µg of chromatin was incubated with rabbit anti-Sox2 (7-10 µg; EMD Millipore #AB5603). For ChIP-H35HT, 10-15 µg chromatin was incubated with rabbit anti-H3K4me3Q5ser (3 µg; EMD Millipore #ABE2580). The next day, lysates were incubated with Protein A/G magnetic beads (Thermo #88802) for 5-6 h at 4°C, followed by washing with Tris-SDS-EDTA-I buffer (0.1% SDS, 1% Triton X-100, 2 mM EDTA, 150 mM NaCl, 20 mM Tris-HCl; pH 8.0), Tris-SDS-EDTA-II buffer (TSEI buffer with 500 mM NaCl), LiCl buffer (250 mM LiCl, 1% Nonidet P-40, 1% sodium deoxycholate, 1 mM EDTA, 10 mM Tris-HCl pH 8.0), and Tris-EDTA buffer (10 mM Tris-HCl pH 8.0, 1 mM EDTA). To release DNA fragments, samples were incubated in freshly prepared elution buffer (1% SDS, 0.1 M $NaHCO_3$) for 20 minutes at 65°C twice. Elutions were treated with proteinase K (0.4 mg/ml: ThermoFisher #AM2546) and NaCl (0.125 M) overnight at 65°C for reverse crosslinking. Subsequently, ChIP-DNA was purified using a PCR purification kit (Qiagen #28104) and quantified using the Quant-iT dsDNA Assay kit. For ChIP-PCR, additional control samples were prepared, wherein sonicated lysates were incubated with rabbit anti-IgG (R&D Systems #AB-105-C). Subsequently, purified ChIP-DNA were analyzed in PCR reactions using AccuPrime Pfx DNA polymerase (ThermoFisher #12344-032) that were carried out at 95°C for 5 min; 40 cycles of 95°C for 30 s, 55°C for 1 min and 68°C for 30 s; followed by 68°C for 2 min. *Slc22a3* primers used for ChIP-PCR was designed at Sox9 binding site at *Slc22a3* promoter (forward 5'-CTGTCCCTCTGTCCCATTGT-

3', reverse 5'- TTCCAGGATCACCCAGACTC-3'). For ChIP-Seq experiments, 10-12 ng of ChIP-DNA was used for library preparation as described below.

ChIP-Seq library preparation, sequencing, and bioinformatic analysis.
ChIP libraries were prepared using the TruSeq ChIP Library Preparation Kit (Illumina #IP-202-1012), according to the manufacturer's instructions. Libraries ranging from 250 to 350 bp were extracted from gel incisions using the QIAquick Gel Extraction Kit (Qiagen #28706), PCR amplified, and purified using AMPure XP beads (Beckman Coulter Life Science #A63882). The quality of the resulting libraries was analyzed on the Standard Sensitivity NGS Fragment Analysis Kit (Agilent #DNF-473-0500) on a 12-capillary Fragment Analyzer. Libraries were quantified using the Quant-iT dsDNA assay kit (ThermoFisher #Q33120), and equal concentrations (2 nM) of libraries were pooled and subjected to single-end (read 1: 150) sequencing of ~60-80 million reads per sample using the High Output v2 kit (Illumina #FC-404-2002) on a NextSeq550 following the manufacturer's instructions.
Sequencing files from each flow cell lane were downloaded, and the resulting fastq files were merged. Quality control was performed using fastQC (v0.11.17) and MultiQC (v0.9). Reads were mapped to the mouse genome mm10 assembly using bowtie2 (v 2.2.6) (*62*). Using the HOMER (v4.10) software suite (*63*), bedgraph files and tag directories were made. The findPeaks command in factor (for Sox9, Sox2) or histone (for H3-5HT) mode was used to filter ChIP peaks enriched over input control. Annotation of enriched peaks was performed using annotatePeaks with mm10 assembly. Integrated Genome Browser–compatible files were made using samtools (v1.9), sort and index, deepTools (v3.2.0), and bamCompare (*64*, *65*). Overlapping peaks, unique peaks, and differentially bound peaks were obtained using mergePeaks or getDifferentialPeaks, and peaks were visualized using computeMatrix and plotHeatmap. Motif analysis for transcription factors was performed using findMotifsGenome.pl at 1000 bp from peak center. GOs were determined by submitting genes associated with ChIP peaks at Enrichr, and significant GO terms ($p < 0.01$) were selected for visualization using ggplot.

Immunoprecipitation followed by mass spectrometry
We collected olfactory bulbs from 6-8 mice for each experimental cohort that were immediately snap frozen. Tissues were thawed, pellet homogenized, and nuclear lysates extracted using NE-PER Nuclear and Cytoplasmic Extraction Reagents (ThermoFisher #78833) according to manufacturer instructions. Lysates were ultracentrifuged at 200,000 *g* for 20 min at 4°C and the supernatant (3-5 mg total protein) used for immunoprecipitation with anti-Sox9 (5 µg, Abcam #ab5535), which were incubated with lysates for 1 hour at 4°C, followed by incubation with protein A Sepharose slurry (GE Healthcare Life Sciences) for another 1 hour at 4°C. The beads were collected by centrifugation at 1000 *g* for 1 min, washed with NETN buffer (50 mM Tris pH 7.3, 1 mM EDTA, 0.5% NP-40) multiple times, and heated for 10 min at 90°C with 20 ml of 2X SDS loading dye to elute bound proteins. Negative control samples were prepared using the same methods without addition of anti-Sox9 antibody. The immunoprecipitated samples were resolved on NuPAGE 10% Bis-Tris Gel (Life Technologies) and the gel pieces were processed for in-gel digestion using Trypsin enzyme (GenDEPOT #T9600). The tryptic peptides were analyzed on nano-LC 1200 system (Thermo Fisher Scientific, San Jose, CA) coupled to Orbitrap Fusion Lumos (Thermo Fisher Scientific, San Jose, CA) mass spectrometer. The MS/MS spectra was searched using Mascot algorithm (Mascot 2.4, Matrix Science) against the mouse NCBI refseq protein database in the Proteome Discoverer (PD1.4, Thermo Fisher) interface. The precursor mass tolerance was confined to 20 ppm, fragment mass tolerance of 0.5dalton, maximum of two missed cleavage was allowed. Dynamic modification of oxidation on methionine, protein N-terminal Acetylation, destreak on cysteine and phosphorylation on serine,

threonine and tyrosine was allowed. The assigned peptides were filtered at 5% FDR using the percolator q-value. The protein quantification was performed using the iBAQ approach.

Western blot
Whole cell lysates were prepared in RIPA lysis buffer, run on a 10% sodium-dodecyl sulfate polyacrylamide gel, followed by wet transfer to nitrocellulose membrane at 400 mA for 45 minutes. The membrane was blocked by 5% milk in Tris-buffered saline with Tween20 (TBST), followed by incubation overnight at 4°C. The following primary antibodies were used: rabbit anti-Sox9 (1:500 dilution, Abcam #ab5535), rabbit anti-Sox2 (1:500, EMD Millipore #AB5603), rabbit anti-Fos (1:500, Cell Signaling #2250S), mouse anti-Gapdh (1:500 dilution, EMD Millipore #MAB374). The next day, membranes were washed three times with TBST, incubated at room temperature for 1 hour in horseradish peroxidase–conjugated IgG at 1:2000 dilution in 5% milk, washed again three times with TBST, and developed using luminol reagent (Santa Cruz Biotechnology #sc2048). For western blot after immunoprecipitation, nuclear lysates were prepared as described above for sample preparation for mass spectrometry. The following primary antibodies were used: rabbit anti-Sox9 (1:500 dilution, Abcam #ab5535) and rabbit anti-Sox2 (1:500, EMD Millipore #AB5603). Subsequent pull-down was performed by adding protein A agarose beads (ThermoFisher #15918-014) for an additional 5 hours at 4°C. The beads were collected, washed, and boiled in 2× SDS gel loading dye to elute bound proteins and run for western blot.

Olfactory behavior
Behavioral assays for odor detection and odor discrimination were performed as describe previously (*1*). Odor detection was performed by olfactory habituation/dishabituation in a 3-chamber place preference assay. Mice were acclimated and familiarized to the testing chamber for 4 minutes each day for three days prior to testing. Mice were first habituated to outer chambers containing mineral oil and the middle chamber serving as a neutral barrier for three times, following which mineral oil in one of the outer chambers was swapped with a novel odorant (R)-limonene in an increasing concentration series ($10^{-6}$ – $10^{-3}$ v/v dilution). Odor detection was calculated based on the time spent investigating the odorant containing chamber in comparison to mineral oil chamber and expressed as zone preference index. To assay for differences in odor discrimination, a similar experimental paradigm was used, wherein a different but structurally similar odorant (S)-limonene was introduced, while mineral oil was replaced with (R)-limonene. Mice were first habituated to a high concentration of (R)-limonene ($10^{-3}$ v/v dilution) in both outer chambers before introduction of (S)-limonene ($10^{-3}$ v/v dilution) in one of the chambers. Videos were captured with a Logitech HD 1080p camera. Preference for an odor was determined based on time spent in each chamber and calculated as zone preference index using MATLAB software with the Optimouse plug-in for analysis of mouse positions (*66*).

Imaging and Sholl analysis
Fluorescent images for morphological evaluation were acquired using a Zeiss LSM 980 confocal microscope with 63× oil immersion objective with frame size at 1024 × 1024 and bit depth at 12. Serial images at z axis were taken at an optical step of 0.5 mm, with overall z axis range encompassing the whole section. Images were imported to Imaris Bitplane software and only astrocytes with their soma between the z-axis range were chosen for further analysis. We performed 3D surface rendering using the Imaris Surface module, and color coded the reconstructed surface images based on the surface area of each astrocyte. Morphological analysis was performed using the Imaris Filament module. Astrocyte branches and processes were outlined by Autopath with starting point set at 8 mm and seed point set at 0.7 mm, and statistical outputs including "filament number Sholl intersections" were extracted and plotted. To

measure the shortest distance between Ezrin and PSD95, figures were converted to the Imaris proprietary format (,ims) in IMARIS (OXFORD instruments, 9.7.0).

Two-photon GCaMP6 calcium imaging and analysis

Animals were anesthetized with isoflurane and isolated brains were submerged in ice-cold ACSF solution (130 mM NaCl, 24mM $NaHCO_3$, 1.25 mM $NaH_2PO_4$, 3.5 mM KCl, 1.5 mM $CaCl_2$, 1.5 mM $MgCl_2$, and 10 mM D(+)-glucose, pH 7.4). 300 mm olfactory bulb slices were cut using a vibratome (DSK Linear Slicer, Kyoto, Japan) oxygenated in ACSF. Slices were then recovered in oxygenated ACSF for 15 minutes and allowed to acclimate to room temperature for at least 30 minutes before imaging. We recorded calcium activity using a two-photon resonant microscope (LSM 7MP, Zeiss) equipped with a Coherent Chameleon Ultra (II) Ti-sapphire laser tuned to 900 nm and a 20×, 1.0 NA Zeiss objective. Calcium activity was typically sampled at ~1 Hz. Optical signals were recorded for 5 minutes per trial at 1024 x 1024 pixel resolution. We recorded data from astrocytes at depths of 30 μm below the surface. All multiphoton imaging experiments were performed within 2-4 hours of slicing. For serotonin induced calcium imaging, optical signals were recorded after slices were bathed in 500 nM tetrodotoxin (TTX) for 5 minutes. After 200s of recording under TTX treatment, brain slices were bathed in 50 μM serotonin (ThermoFisher #AAB2126303) and recorded for an additional 300s. For glutamate (Sigma #G1251) induced calcium imaging, slices were bathed in 300 μM instead of serotonin. Image analysis of spontaneous or induced $Ca^{2+}$ were quantified using GECIquant algorithm with ImageJ software, wherein detection of ROI for soma was performed in a semi-automated manner as described in a previous study (*67*, *68*). After thresholding from temporally projected stack images with a maximum intensity projection, a polygon selection was manually drawn around the approximate astrocyte territory of interest, and the selection was added to the ImageJ ROI manager. The area criterion was 30 μm to infinity for soma within the GECIquant ROI detection function. Intensity values for each ROI were extracted in ImageJ and converted to dF/F values. For each ROI, basal F was determined during 40 s periods with no fluctuations. Clampfit 10.7 software was used to detect and measure amplitude and frequency values for the somatic and microdomain transients. We counted the response following with these criteria: amplitude (> 0.5 dF/F), pre - trigger time (3 ms), and minimum duration (5 ms). Data points were plotted on Prism software.

Slice recording for EPSC, IPSC and tonic GABA

Animals were anesthetized with isoflurane and isolated brains were submerged in ice-cold ACSF solution (130 mM NaCl, 24mM $NaHCO_3$, 1.25 mM $NaH_2PO_4$, 3.5 mM KCl, 1.5 mM $CaCl_2$, 1.5 mM $MgCl_2$, and 10 mM D(+)-glucose, pH 7.4). 300 μm slices were cut using a vibratome (DSK Linear Slicer, Kyoto, Japan) oxygenated in ACSF at room temperature for 1 h, and then acclimated at room temperature with continuous perfusion with ASCF solution (2 ml/min). Slices were placed in recording chamber and target cells were identified via upright Olympus microscope with a 60× water immersion objective with infrared differential interference contrast optics. Whole cell recording was performed with pCLAMP10 and MultiClamp 700B amplifier (Axon Instrument, Molecular Devices) at room temperature from olfactory bulb granule cells. The holding potential was -60 mV. Pipette resistance was typically 5-8 MΩ. The pipette was filled with an internal solution (in mM): 135 CsMeSO4, 8 NaCl, 10 HEPES, 0.25 EGTA, 1 Mg-ATP, 0.25 Na2-GTP, 30 QX-314, pH adjusted to 7.2 with CsOH (278-285 mOsmol) for EPSC measurement , 135 CsCl, 4 NaCl, 0.5 $CaCl_2$, 10 HEPES, 5 EGTA, 2 Mg-ATP, 0.5 Na2-GTP, 30 QX-314, pH adjusted to 7.2 with CsOH (278-285 mOsmol) for inhibitory postsynaptic currents (IPSCs) and tonic current measurement. IPSC and tonic current were measured in the presence of ionotropic glutamate receptor antagonists, APV (50 μM, Tocris), and CNQX (20 μM, Tocris). Electrical signals were digitized and sampled at 50 μs intervals with Digidata 1550B and

Multiclamp 700B amplifier (Molecular Devices, CA, USA) using pCLAMP 10.7 software. Data were filtered at 2 kHz. The recorded current was analyzed with ClampFit 10.7 software. Data points were plotted on Prism software.

Statistical analysis

Sample sizes and statistical tests are provided in all figure legends. Summary data of all mean, SEM, p-values, sample sizes and statistical methods used are provided in Supp Table S3. Data were tested for normality using the Shapiro–Wilk tests and for homogeneity of variance using the Levene test. Parametric tests were used for normally distributed data sets, and for data with a small sample size (n= 3). For comparison of two groups, unpaired or paired Student's t-test were used, and for comparison of three groups, one-way ANOVA followed by Tukey's tests were used. For multiple comparisons, two-way ANOVA with Sidak's corrected were used. When data did not follow a normal distribution, non-parametric Wilcoxon rank sum tests were applied. Significant differences are denoted by asterisks in associated graphs. Data in bar graphs presented as ± SEM. Box plots were generated using ggsignif (v0.6.0) with ggplot2 (v3.3.2). Levels of statistical significance are indicated as follows: *$p < 0.05$, **$p < 0.01$, ***$p < 0.001$, **** $p < 0.0001$.

### Supporting Figures:

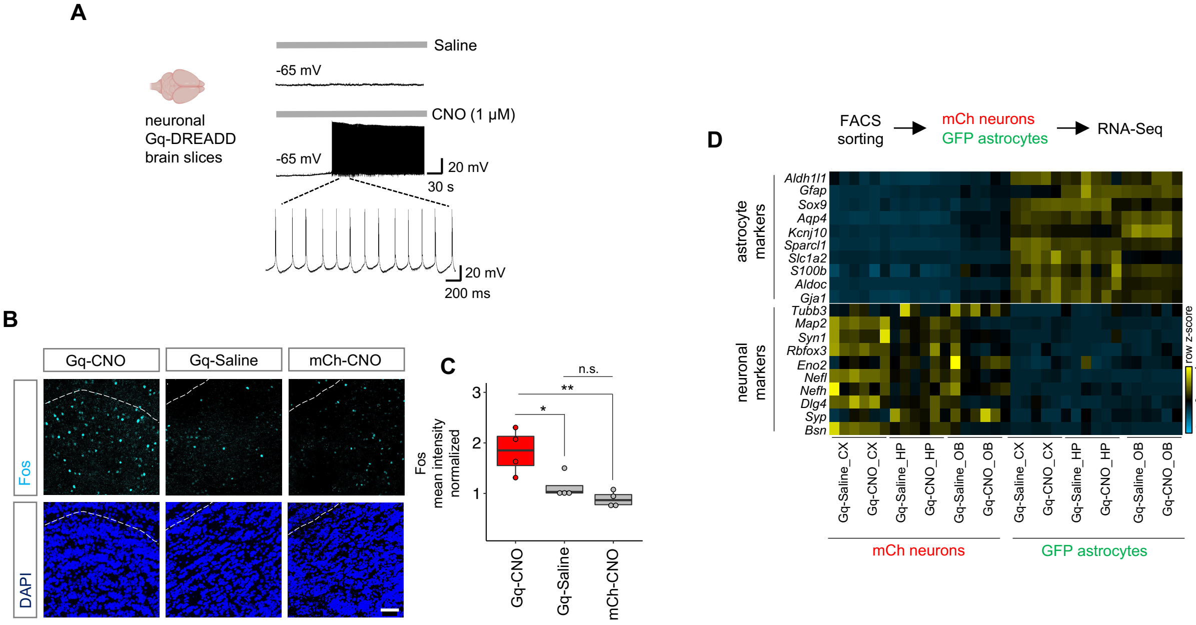


**Supp Fig. S1**
(**A**) Enhanced neuronal firing was observed after CNO application to brain slices expressing neuronal Gq-DREADD, in comparison to saline application. (**B**) Expression of Fos in CNO (Gq-CNO) and saline (Gq-Saline) treated brains from Gq-DREADD mice, and control of CNO treatment (mCh-CNO) in empty viral vector expressing mice. Sections shown are from the OB. Scale bar: 50 μm. (**C**) Box plot depicting quantification of Fos mean fluorescence intensity in Gq-CNO, Gq-Saline, and mCh-CNO. **p* = 0.0336 (Gq-CNO vs. Gq-Saline); **p = 0.0069 (Gq-CNO vs. mCh-CNO); *p*= 0.1385 (Gq-Saline vs. mCh-CNO); one-way ANOVA with Tukey's test on n= 4 mice/cohort. (**D**) Expression heatmap of neuronal and astrocyte markers from RNA-Seq data of FACS sorted neurons and astrocytes in Gq-CNO and Gq-Saline samples over different brain regions of cortex (CX), hippocampus (HP) and olfactory bulb (OB). Note enrichment of neuronal and astrocyte markers in sorted mCh neurons and GFP astrocytes, respectively (n= 3 each cohort).

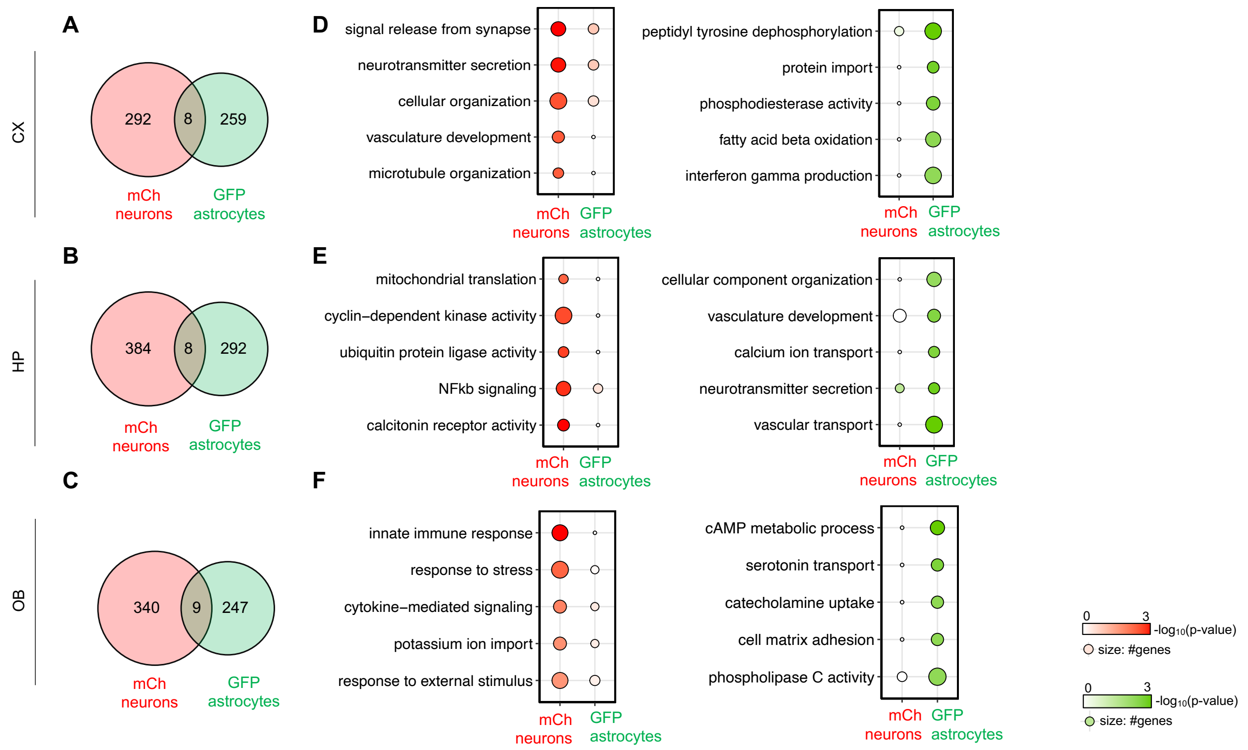


**Supp Fig. S2**

(**A-C**) Number of differentially expressed genes (DEGs) in Gq-CNO vs. Gq-Saline treatment in Gq-DREADD expressing mice in FACS sorted mCh neurons and GFP astrocytes across CX, HP and OB (n= 3 each cohort, $p$< 0.05, $\log_2$fold-change >1). (**D-F**) Enriched gene ontology (GO) terms associated with cell type specific and region-specific DEGs.

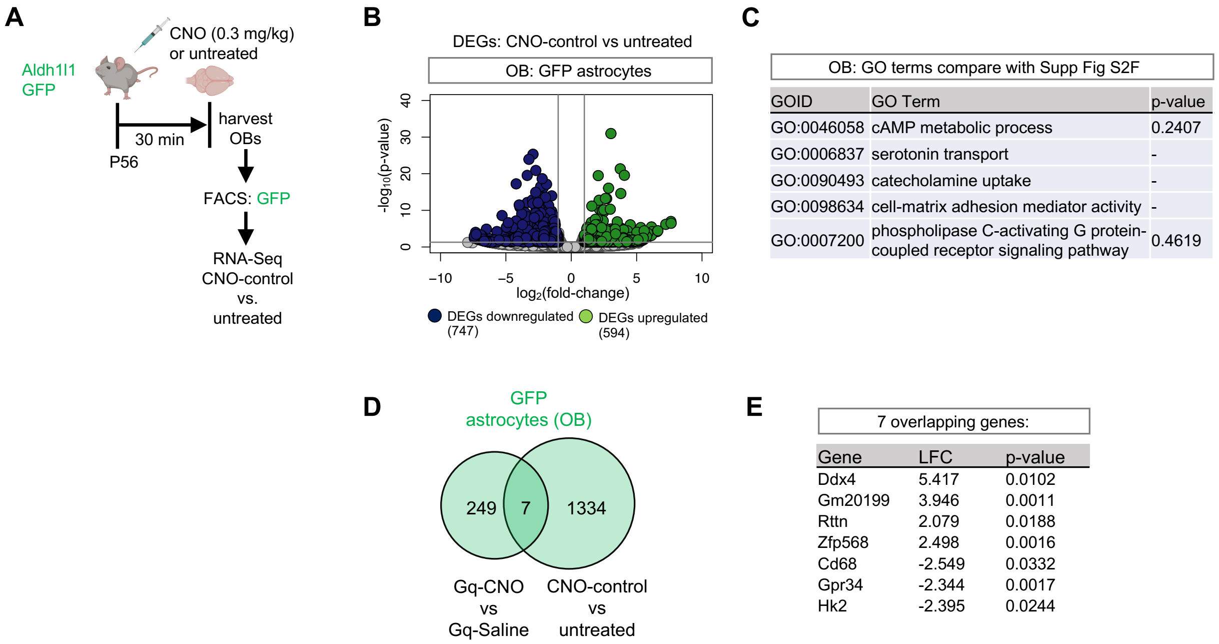


**Supp Fig. S3**

(**A**) Schematic of control experiment comparing CNO treatment (CNO-control) in comparison to untreated Aldh1l1-GFP mice (untreated). (**B**) Volcano plots depicting RNA-Seq from FACS sorted GFP astrocytes comparing CNO-control with untreated. (**C**) Level of significance of the same GO terms enriched in Gq-CNO vs Gq-Saline astrocytic DEGs (shown in Figure S2F) in CNO-control vs. untreated DEGs. (**D-E**) Venn diagram and gene names the 7 overlapping genes between DEGs from Gq-CNO vs. Gq-Saline (shown in Figure 1E; OB) and CNO-control vs. untreated.

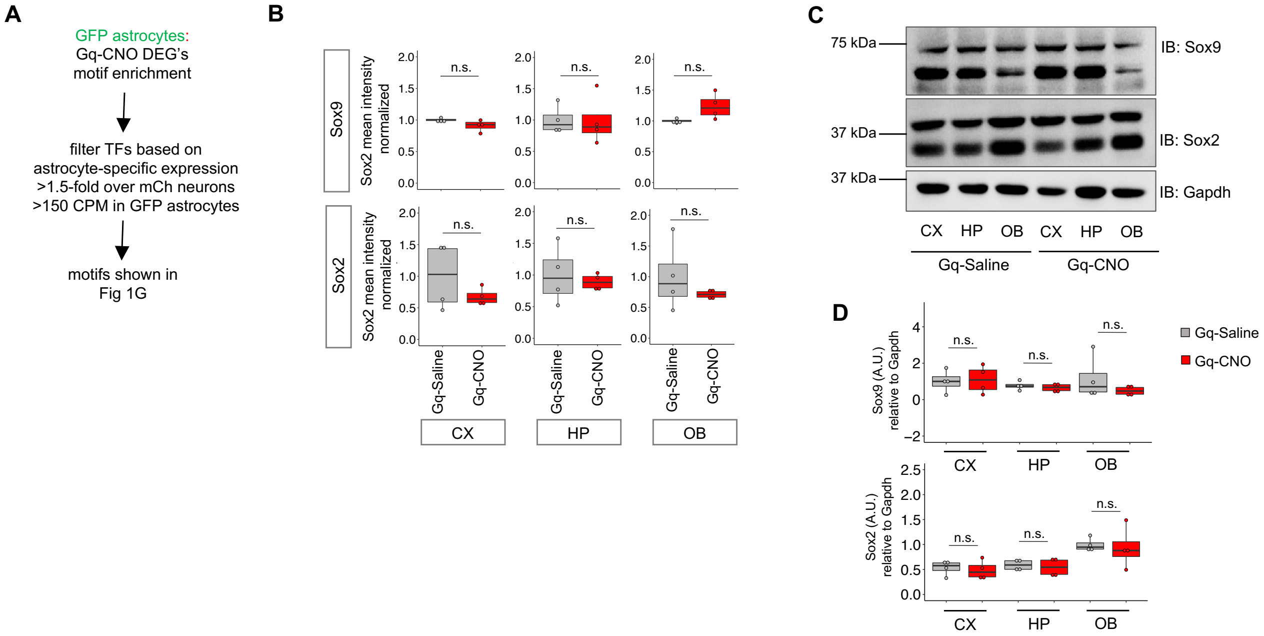


**Supp Fig. S4**

(**A**) Schematic of motif enrichment analysis in GFP astrocytes. (**B**) Sox9 and Sox2 protein expression obtained from immunostaining analyses in Gq-CNO vs. Gq-Saline across CX, HP, and OB. Note no significant change in expression after Gq-DREADD neuronal activation. Sox9: CX, $p$= 0.1049; HP, $p$= 0.9707; OB, $p$= 0.0591; unpaired Student's two-tailed t-test or Wilcoxon rank sum test on n= 4 mice/cohort. Sox2: $p$= 0.6857; HP, $p$= 0.6618; OB, $p$= 0.4857; unpaired Student's two-tailed t-test or Wilcoxon rank sum test on n= 4 mice/cohort. (**C**) Sox9 and Sox2 protein expression obtained from western blot quantification in Gq-CNO vs. Gq-Saline treatments across CX, HP, and OB. Note no significant change in expression after Gq-DREADD neuronal activation. Loading control: GAPDH. Sox9: CX, $p$= 0.8524; HP, $p$= 0.4809; OB, $p$= 0.3256; unpaired Student's two-tailed t-test or Wilcoxon rank sum test on n= 4 mice/cohort. Sox2: $p$= 0.7068; HP, $p$= 0.6724; OB, $p$= 0.7967; unpaired Student's two-tailed t-test or Wilcoxon rank sum test on n= 4 mice/cohort.

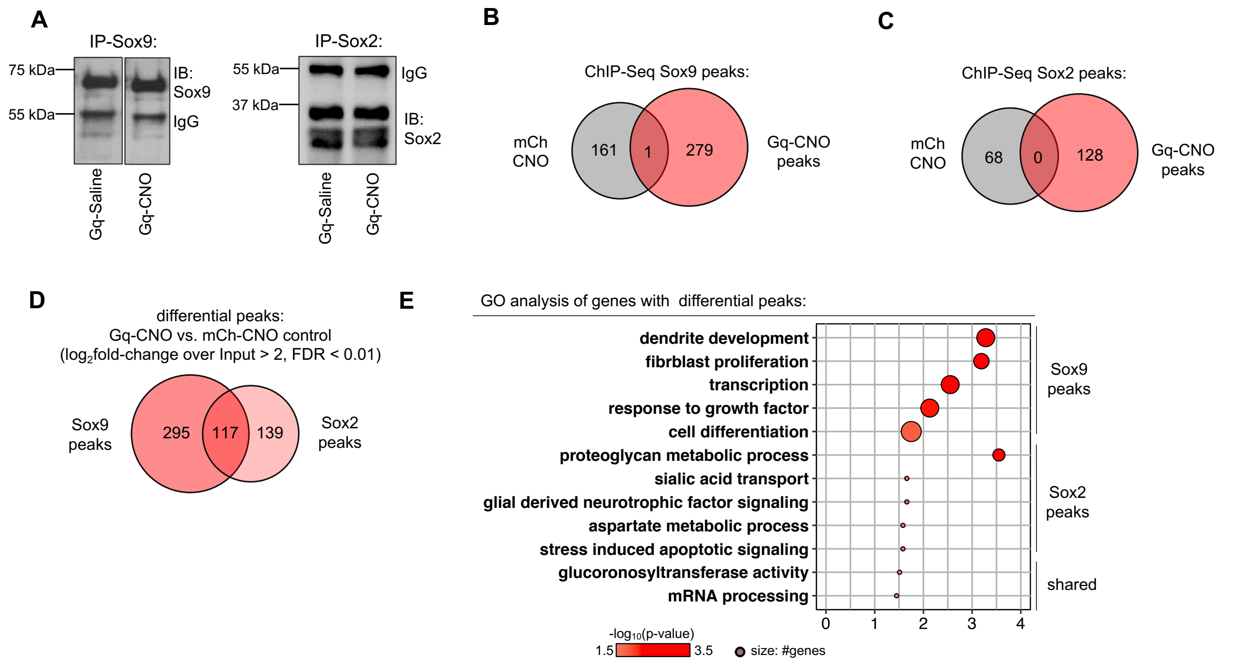


**Supp Fig. S5**

(**A**) Similar levels of Sox9 and Sox2 are immunoprecipitated in Gq-CNO vs Gq-Saline controls. (**B-C**) Venn diagrams depicting ChIP-Seq Sox9 and Sox2 peaks in Gq-CNO in comparison to mCh-CNO controls (n= 3/cohort). (**D**) Differential peak analysis between ChIP-Sox9 and ChIP-Sox2 in Gq-CNO brains, and (**E**) GO analysis of genes at peaks obtained from differential peak analysis.

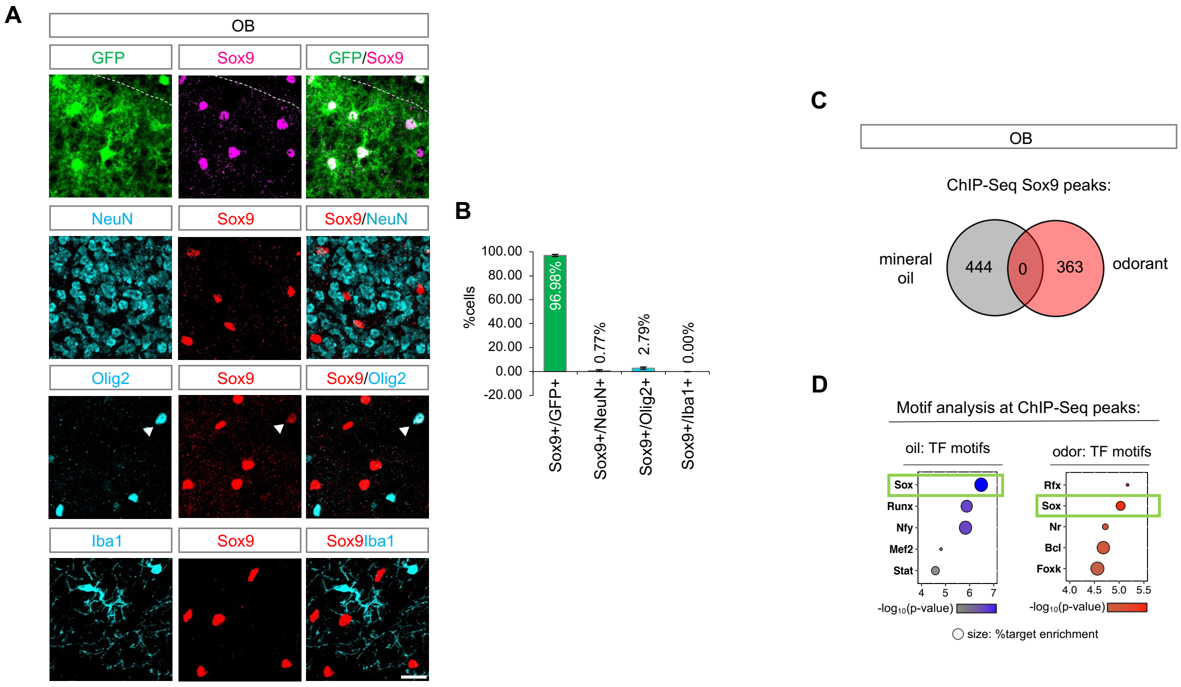


**Supp Fig. S6**

(**A-B**) Representative images and quantification showing Sox9 co-labeling with different cell type markers. Note 96.983 ± 0.873 percent enrichment in Aldh1l1-GFP+ astrocytes, 2.793 ± 1.049 percent detected in oligodendroglial lineage, 0.771 ± 0.770 percent in neurons, and none in microglia (n= 3 mice, 3 sections each). Scale bar: 20 μm. (**B**) Venn diagrams depicting ChIP-Seq Sox9 in OBs from mice after mineral oil and odor treatment and (**C**) motif analysis at identified ChIP-Seq Sox9 peaks in oil and odor treatments (n= 6 each cohort).

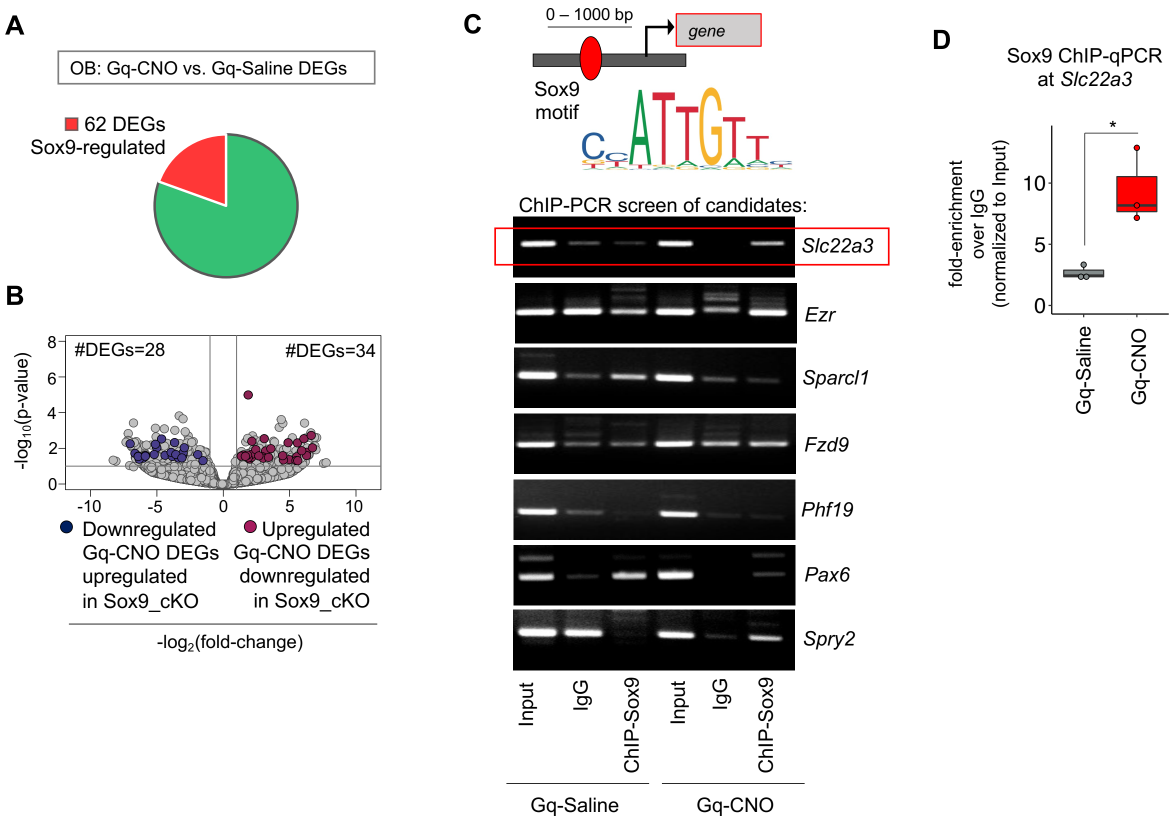


**Supp Fig. S7**

(**A**) Pie-chart and (**B**) volcano plot depicting neuronal activity-dependent OB astrocyte-specific DEGs that are also differentially expressed in Sox9-cKO OB astrocytes (n= 3, *p*< 0.05, $\log_2$fold-change 1). (**C**) Schematic depicting ChIP-PCR screen parameters of candidate genes and ChIP'd DNA fragments from OBs of Gq-CNO vs. Gq-Saline treatments. Anti-IgG and anti- Sox9 antibodies were used for ChIP. PCR primers were designed to include predicted Sox9 binding site at candidate gene promoters. Note Sox9 binding at *Slc22a3* promoter specifically in Gq-CNO OBs. (**D**) Quantitative ChIP-PCR of Sox9 at *Slc22a3* promoter (each data point pool of 2–3 OBs, **p*= 0.0199, unpaired Student's two-tailed t-test on n= 3

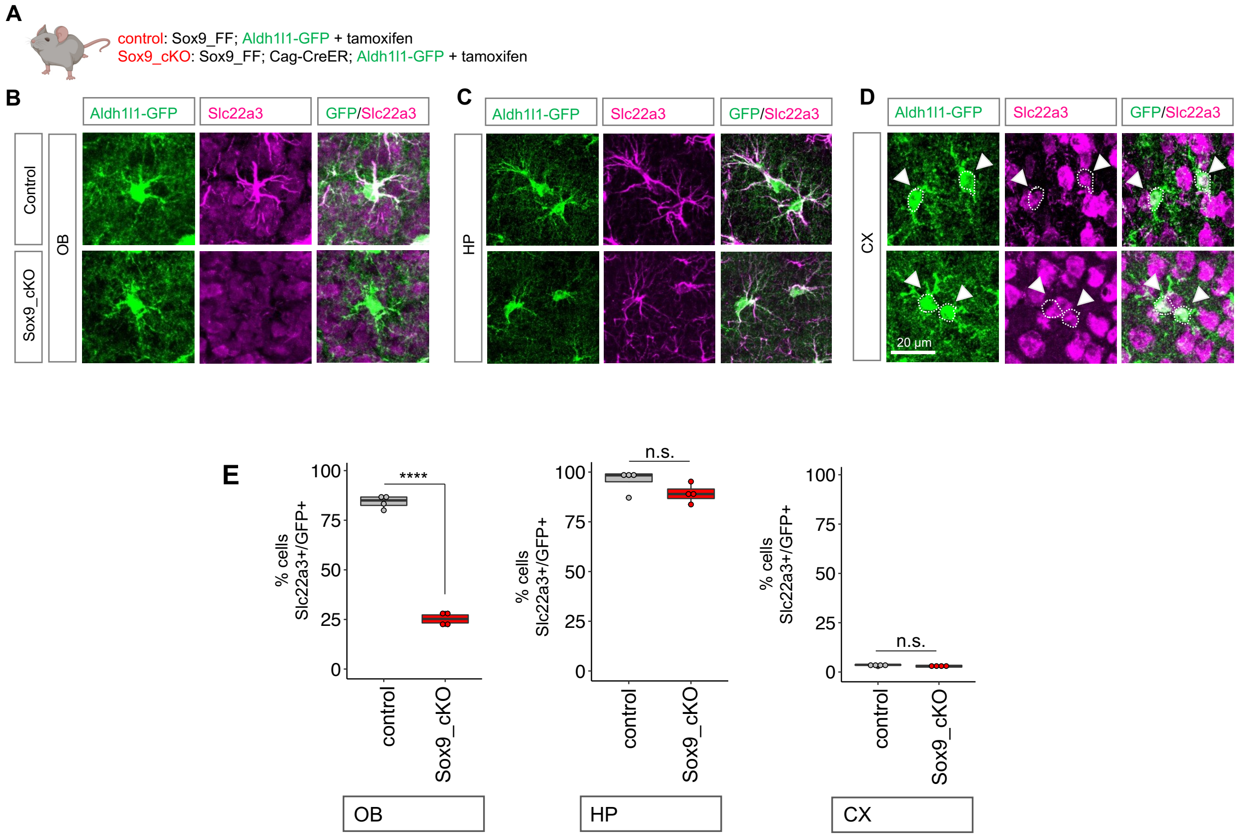


**Supp Fig. S8**

(**A**) Schematic depicting experimental design for Sox9-cKO. (**B-D**) Immunostaining and (**E**) quantification of Slc22a3 in control and Sox9-cKO across OB, HP and CX over Aldh1l1-GFP. (OB: ****$p$= 2.89e-07; HP: $p$= 0.133; CX: $p$= 0.209, unpaired Student's two-tailed t-test on n= 4). Slc22a3 quantification was based on expression levels in soma and astrocyte processes, and very low expression levels were observed in GFP astrocyte processes in the CX.

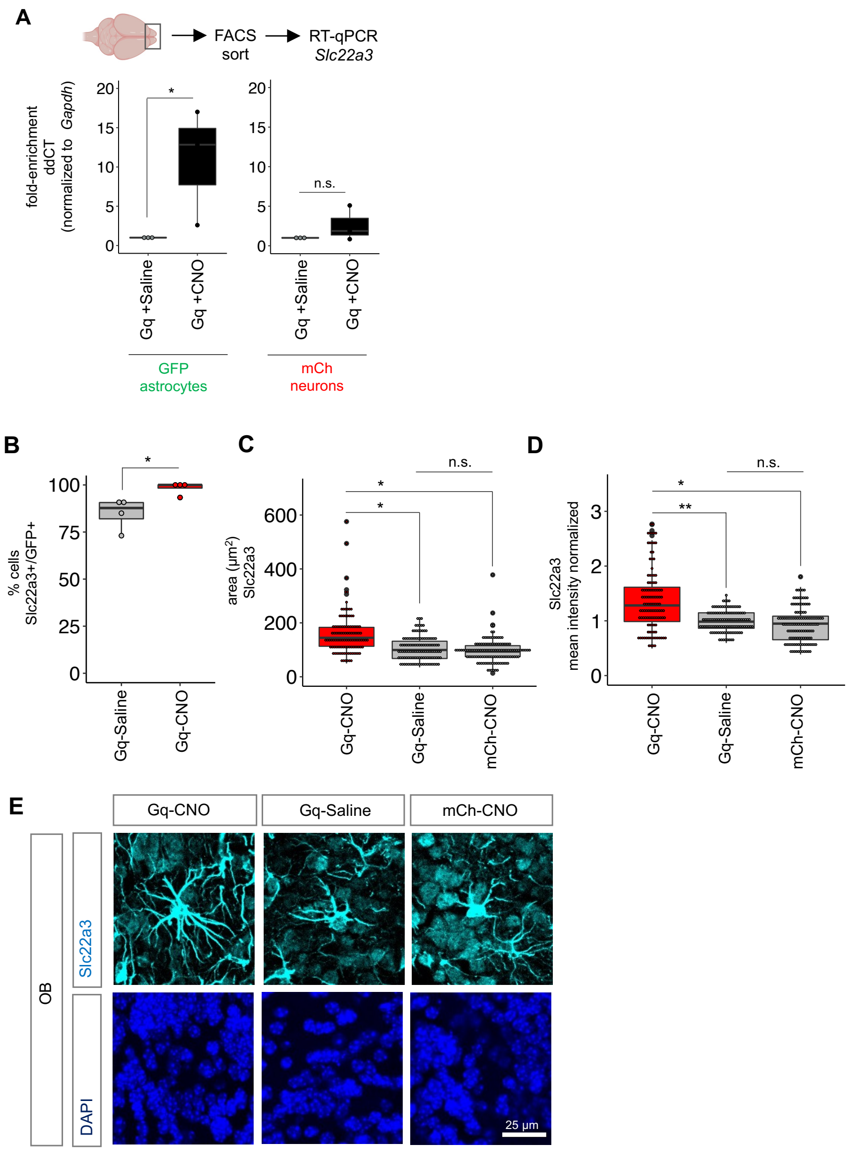


**Supp Fig. S9**

(**A**) RT-qPCR of *Slc22a3* in Gq-CNO vs. Gq-Saline treatments in both cell types of neurons (mCh) and astrocytes (GFP) obtained after FACS sorting (n= 3, GFP astrocytes: **p*= 0.0419; mCh neurons: *p*= 0.1398; unpaired Student's one-tailed t-test). (**B-D**) Quantification and (**E**) representative images of Slc22a3 over GFP astrocytes in Gq-CNO treatment compared to controls. Box plots showing quantification of %cells: **p*= 0.0265 (each data point average of 3 sections, Wilcoxon rank sum test on n= 4 mice/cohort); area: **p*= 0.0423 (Gq-CNO vs. Gq-Saline), **p*= 0.0259 (Gq-CNO vs. mCh-CNO), *p*= 0.5901 (Gq-Saline vs. mCh-CNO) (113-127 cells/cohort, one-way ANOVA with Tukey test on n= 4 mice/cohort); mean intensity: ***p*= 0.0099 (Gq-CNO vs. Gq-Saline), **p*= 0.0162 (Gq-CNO vs. mCh-CNO), *p*= 0.4672 (Gq-Saline vs. mCh-CNO); 115-127 cells/cohort, one-way ANOVA with Tukey test on n=4 mice/cohort.

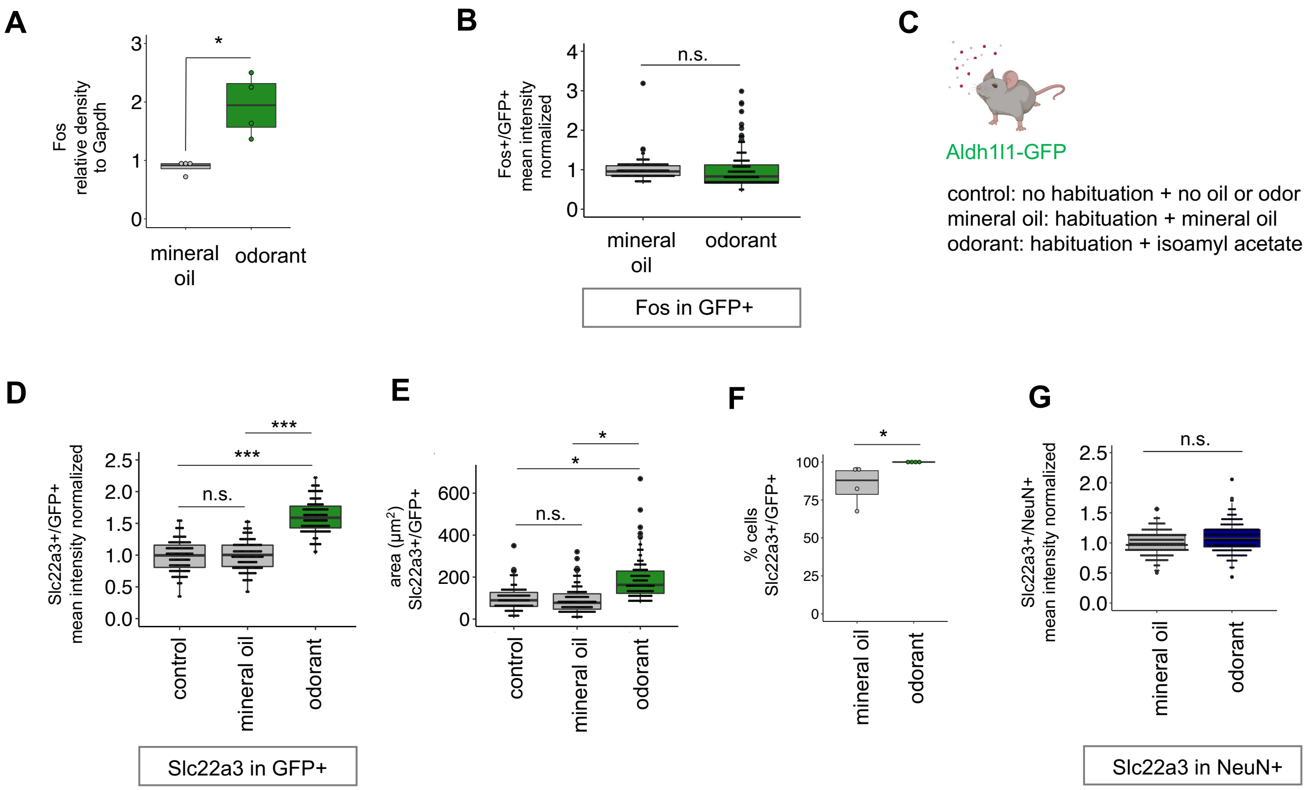


**Supp Fig. S10**

(**A**) Fos protein expression obtained from western blot quantification of OBs from mineral oil vs. odor treatments (*p*= 0.0285, Wilcoxon rank sum test on n= 4 mice/cohort). (**B**) Quantification of mean fluorescence intensity of Fos in Aldh1l1-GFP astrocytes after oil and odorant exposure (128-130 cells/cohort, *p*= 1, Wilcoxon rank sum test on n= 4 mice/cohort). (**C**) Details of experimental and control groups for oil and odor treatment to mice and (**D-F**) box plots showing quantification of Slc22a3 over GFP depicting mean intensity: *p*= 0.8937 (control vs. mineral oil), ****p*= 0.0009 (control vs. odor), ****p*= 0.0004 (oil vs. odor), 118-128 cells/cohort, one-way ANOVA with Tukey test on n= 4 mice/cohort. Area: *p*= 0.6857 (control vs. mineral oil), **p*= 0.0285 (control vs. odor), **p*= 0.0285 (oil vs. odor), 118-128 cells/cohort, one-way ANOVA with Tukey test on n= 4 mice/cohort). Percent cells: **p*= 0.0211 (each data point average of 3 sections, Wilcoxon rank sum test on n= 4 mice/cohort). (**G**) Box plot showing quantification of Slc22a3 over NeuN mean intensity (240-241 cells/cohort, *p*= 0.1184, unpaired Student's two-tailed t-test on n= 4 mice/cohort).

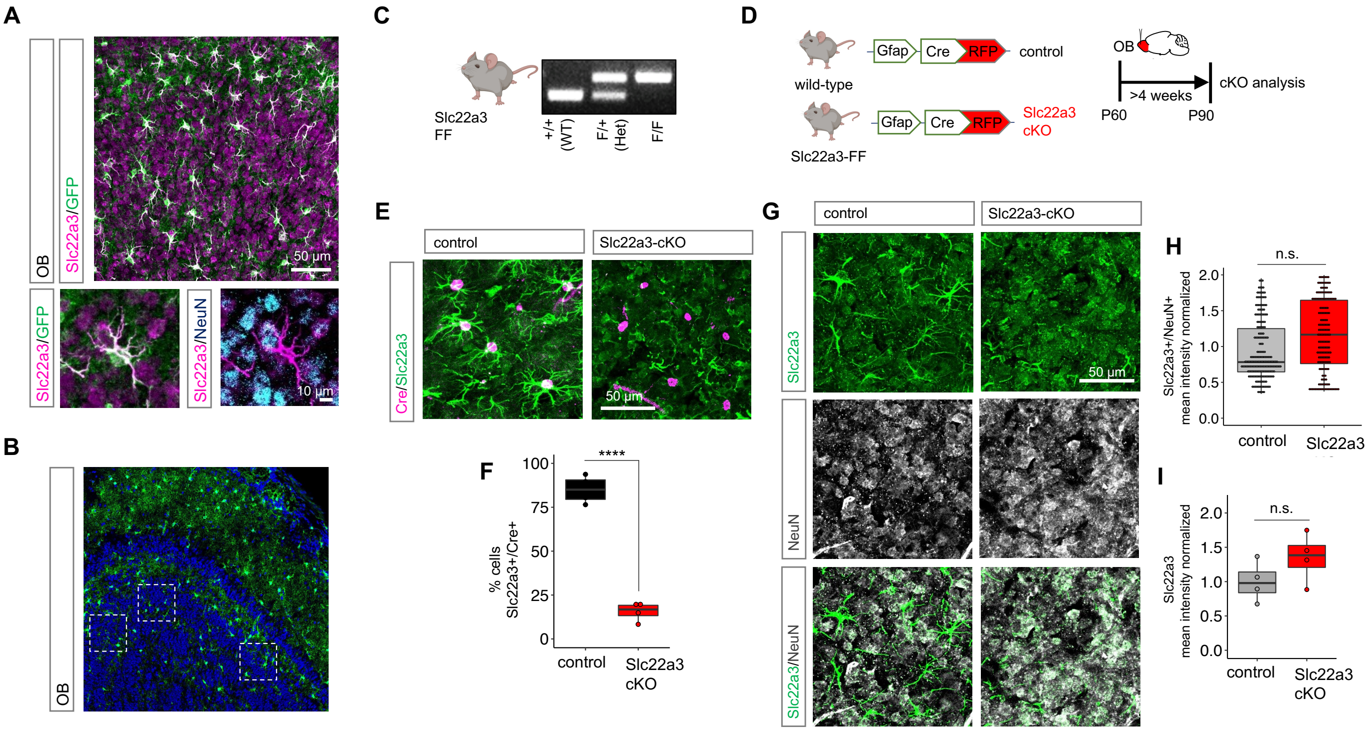


**Supp Fig. S11**

(**A**) Immunostaining of Slc22a3 and co-labeling with Aldh1l1-GFP astrocytes and NeuN+ neurons (**B**) All image analyses performed in this study were obtained from the granular cell layer and sampled regions are shown in white boxes. (**C-D**) Genotyping PCR for Slc22a3-FF in comparison to heterozygous and wild-type controls, and schematic for Slc22a3-cKO vs. control. (**E**) Co- labeling, and (**F**) quantification of Slc22a3 over Cre in control and Slc22a3-cKO (each data point average of 3 sections, ****$p$= 7.03e-06, unpaired Student's two-tailed t-test on n= 4 mice/cohort). (**G**) Co- labeling, and (**H**) quantification of Slc22a3 over NeuN in control and Slc22a3-cKO (215-216 cells/cohort, $p$= 0.2296, unpaired Student's two-tailed t-test on n=4 mice/cohort). (**I**) No change in whole field levels of Slc22a3 was observed between control and Slc22a3-cKO (each data point average of 3-4 sections, $p$= 0.1813, unpaired Student's two-tailed t-test on n=4 mice/cohort).

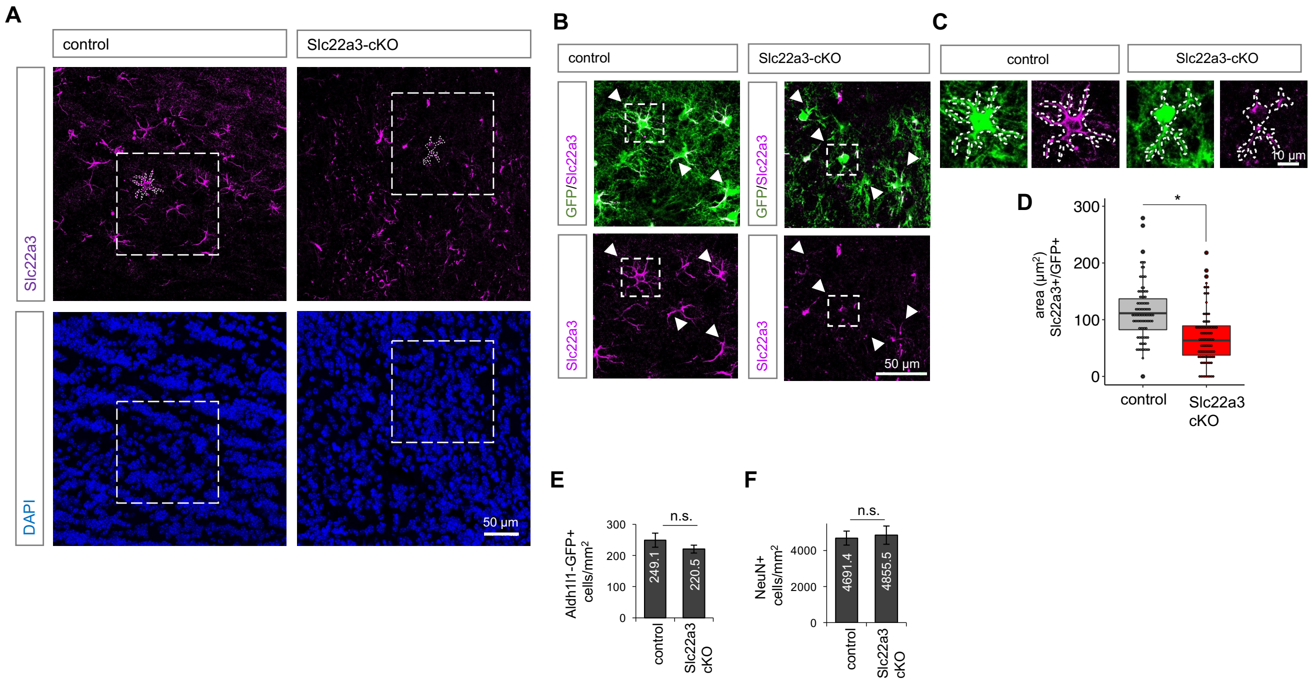


**Supp Fig. S12**

(**A-C**) Zoomed out and merged images of Slc22a3 and GFP immunostaining shown in Fig. 3B. (**D**) Quantification of Slc22a3 area in GFP astrocytes (91-93 cells/cohort, *$p$= 0.0114, unpaired Student's two-tailed t-test on n=4 mice/cohort). (**E-F**) Numbers of GFP+ astrocytes and NeuN+ neurons in OB's of Slc22a3-cKO vs. control (Aldh1l1-GFP: average of 3 sections, $p$= 0.3119, unpaired Student's two-tailed t-test on n=4 mice/cohort; NeuN: average of 3 sections, $p$= 0.8058, Wilcoxon rank sum test on n=4 mice/cohort). Data shown as mean ± SEM.

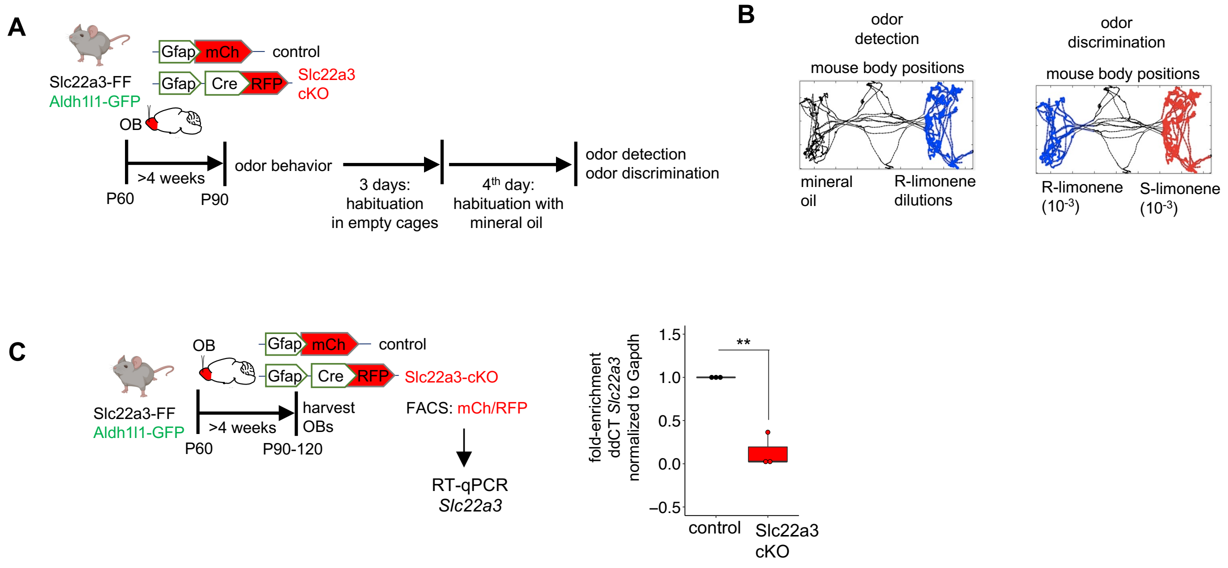


**Supp Fig. S13**
(**A-B**) Schematic illustrating odor behavior protocol and odor detection/odor discrimination behavioral assay design. (**C-D**) Schematic and RT-qPCR quantification of Slc22a3 transcript in Slc22a3-cKO vs. control (*p*= 0.0016, unpaired Student’s two-tailed t-test on n=3 mice/cohort).

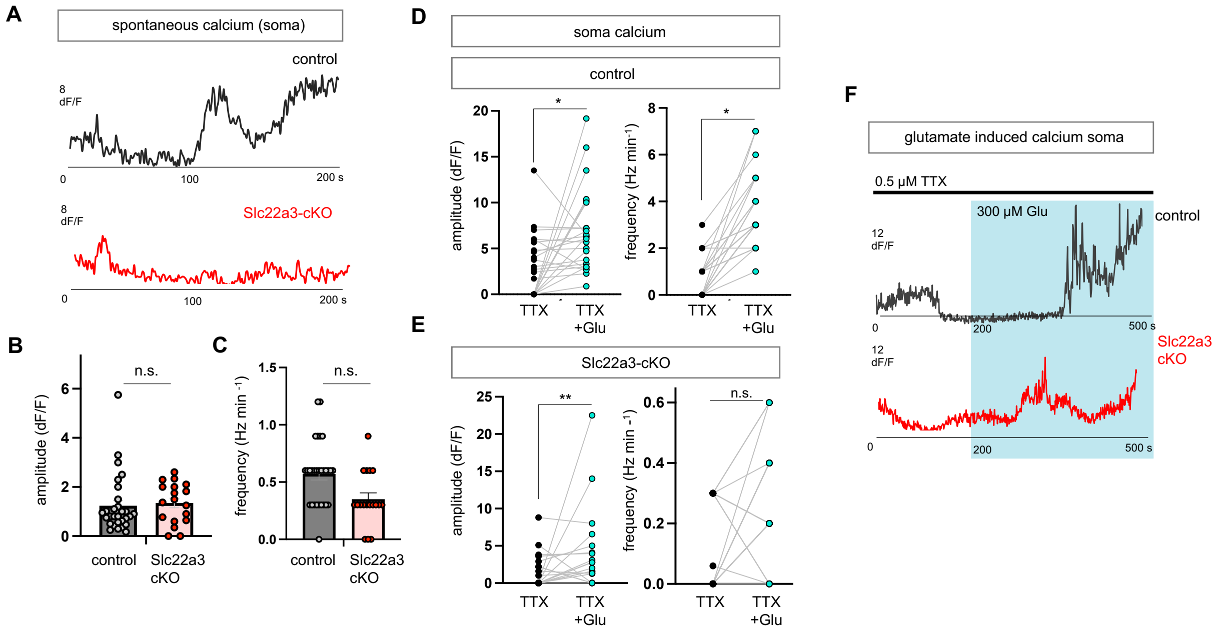


**Supp Fig. S14**

(**A**) Representative traces and (**B**) quantification of spontaneous calcium activity amplitude and frequency using GCaMP6 optical sensor in control and Slc22a3-cKO (18-29 cells, amplitude *p*= 0.6478, frequency *p*= 0.1292, unpaired Student's two-tailed t-test on n=4 mice/cohort). (**D-E**) Quantification of amplitude and frequency from glutamate (Glu) induced calcium activity from astrocyte soma (21-24 cells/cohort, control amplitude **p*= 0.0391; Slc22a3-cKO amplitude ***p*= 0.0089; control frequency **p*= 0.0112; Slc22a3-cKO frequency *p*= 0.0731; paired Student's two-tailed t-test on n= 4 mice/cohort). (**F**) Representative glutamate induced calcium activity traces from astrocyte soma.

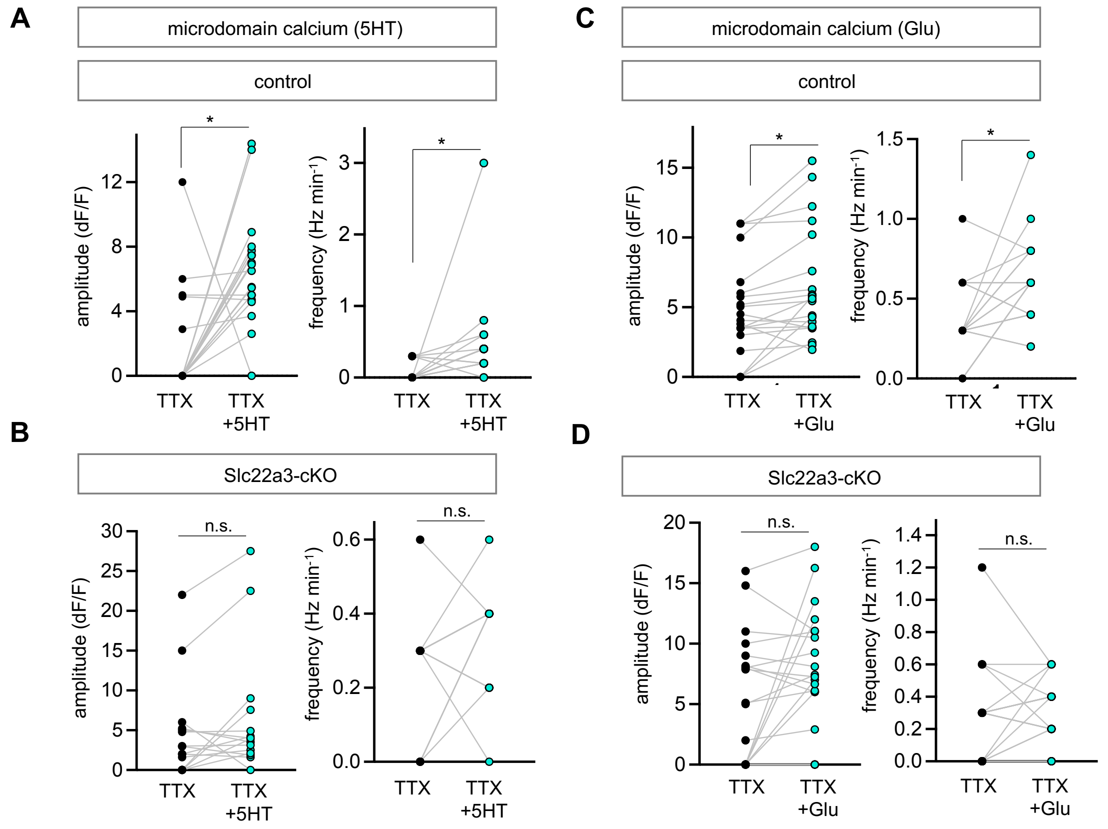


**Supp Fig. S15**
(**A**) Quantification of microdomain calcium activity after application of serotonin (5HT) in control and Slc22a3-cKO OB slices (15-19 cells/cohort, control amplitude *$p$= 0.0466; Slc22a3-cKO amplitude $p$= 0.4889*;* control frequency *$p$= 0.0103; Slc22a3-cKO frequency $p$= 0.3805; paired Student's two-tailed t-test on n= 4 mice/cohort). (**B**) Quantification of microdomain calcium activity after application of glutamate (Glu) in control and Slc22a3-cKO OB slices (20-21 cells/cohort, control amplitude *$p$= 0.0207; Slc22a3-cKO amplitude $p$= 0.0995*;* control frequency *$p$= 0.0203; Slc22a3-cKO frequency $p$= 0.9644; paired Student's two-tailed t-test on n= 4 mice/cohort).

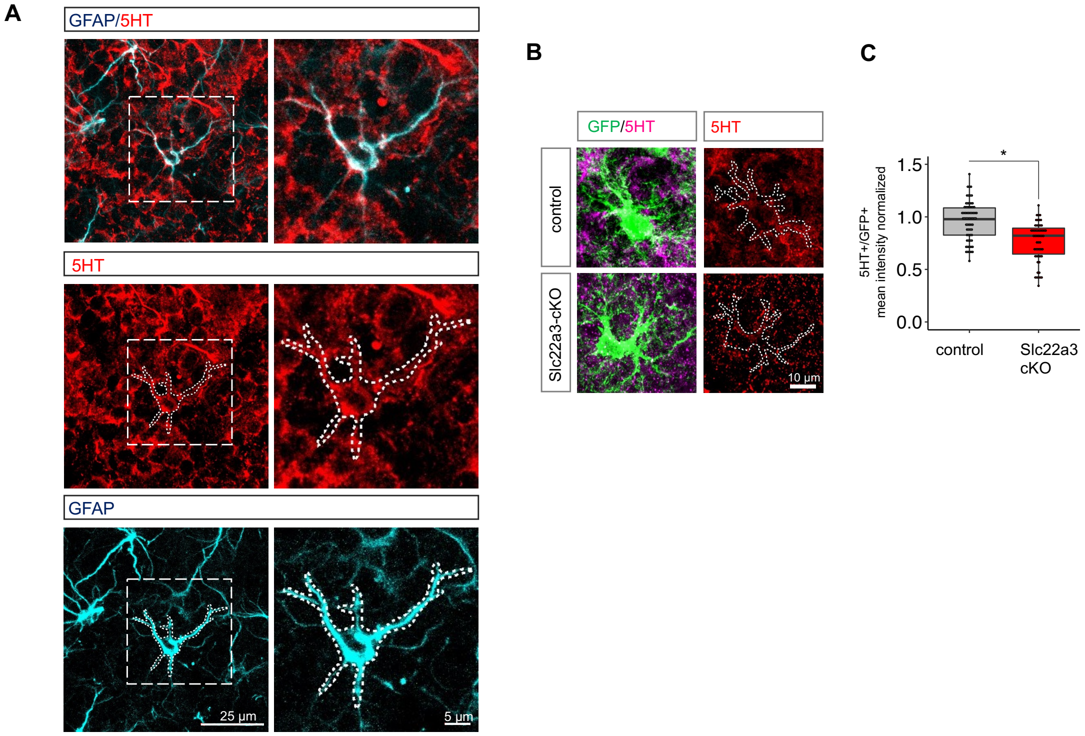


**Supp Fig. S16**
(**A**) Representative immunostaining of serotonin (5HT) in OB of wild-type mice showing presence in astrocytes labeled with marker GFAP. (**B-C**) Representative images and box plot depicting quantification of 5HT in Aldh1l1-GFP astrocytes in Slc22a3-cKO vs. controls (54-67 cells, **p*= 0.0211, unpaired Student's two-tailed t-test on n=4 mice/cohort).

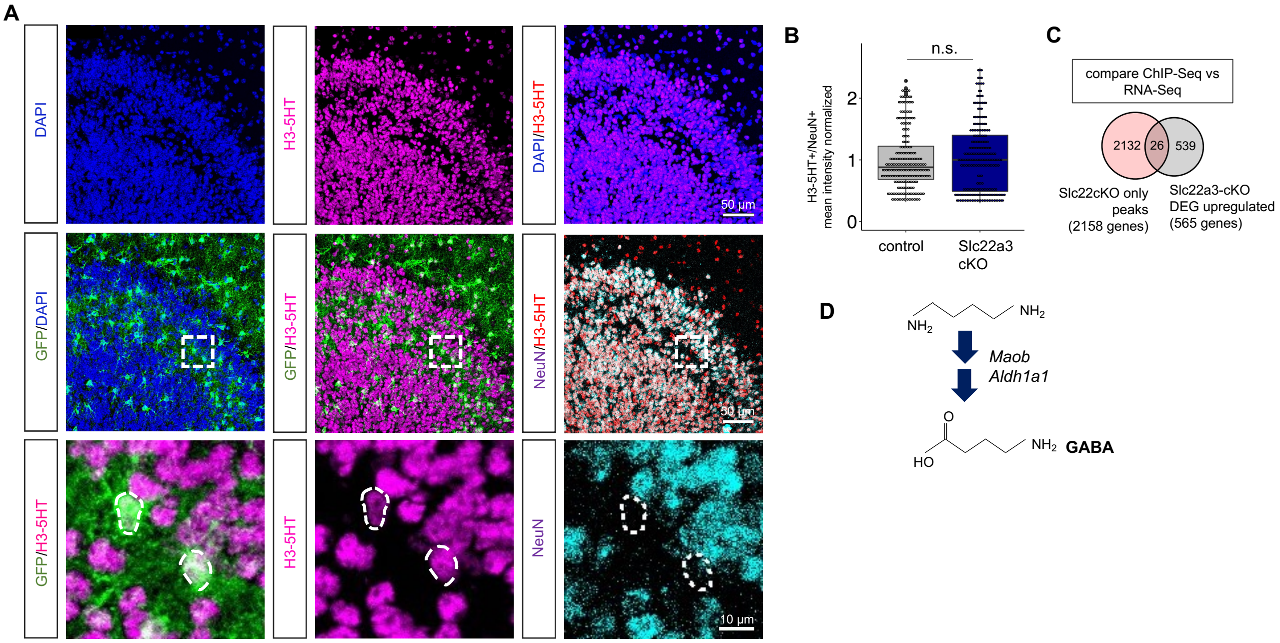


**Supp Fig. S17**
(**A**) Expanded images of H3-5HT and co-labeling with GFP and NeuN immunostaining shown in Fig. 5A. (**B**) Box plot depicting quantification of H3-5HT in neurons (120 cells/cohort, *$p$= 0.9212, unpaired Student's two-tailed t-test on n=4 mice/cohort). (**C**) Venn diagram comparing DEGs upregulated in Slc22a3-cKO and acquiring H3-5HT peaks compared to control. (**D**) Schematic of GABA biosynthesis highlighting enzyme expression levels that were subsequently evaluated.

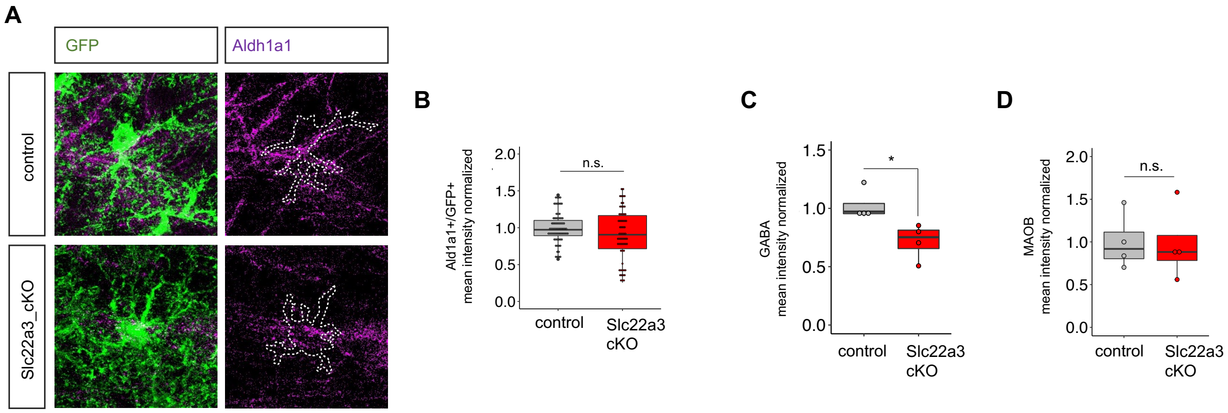


**Supp Fig. S18**

(**A-B**) Representative images and box plot depicting quantification of Aldh1a1 in control vs. Slc22a3-cKO GFP astrocytes in the OB (62-71 cells/cohort, *p*= 0.5807, unpaired Student's two-tailed t-test on n=4 mice/cohort). (**C-D**) Box plot depicting quantification of GABA and MAOB from whole field (each data point average of 3-5 sections, GABA *$p$= 0.0225, MAOB *p*= 0.9345, unpaired Student's two-tailed t-test on n=4 mice/cohort).

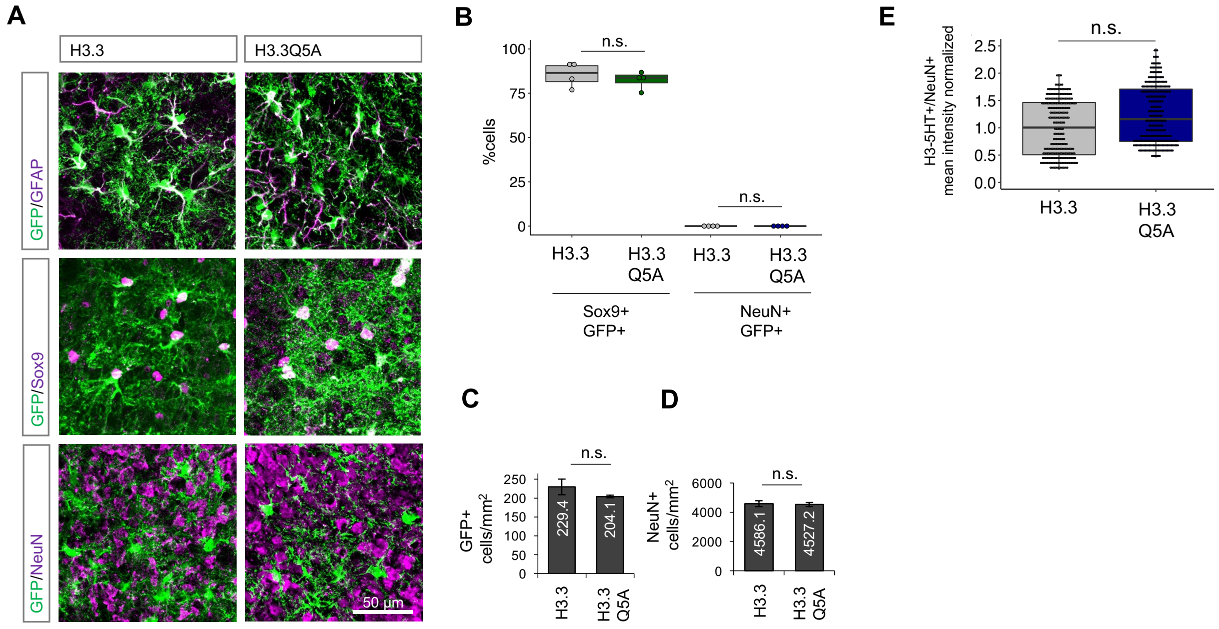


**Supp Fig. S19**

(**A-B**) Immunostaining and quantification of images showing co-labeling of H3.3-GFP and H3.3Q5A-GFP constructs with astrocyte markers Sox9 and GFAP, and absence of co-labeling with NeuN (each data point average of 3 sections, Sox9: $p$= 0.4785, unpaired Student's two-tailed t-test on n=4 mice/cohort). (**C-D**) Numbers of GFP+ astrocytes and NeuN+ neurons in OB's of H3.3Q5A vs. H3.3 (GFP: average of 3 sections, $p$= 0.2733, unpaired Student's two-tailed t-test on n= 4 mice/cohort). Data shown as mean ± SEM. (**E**) Box plot depicting quantification of neuronal H3-5HT in H3.3Q5A vs. H3.3 control OBs (306-332 cells, $p$= 0.4964, unpaired Student's two-tailed t-test on n= 4 mice/cohort).

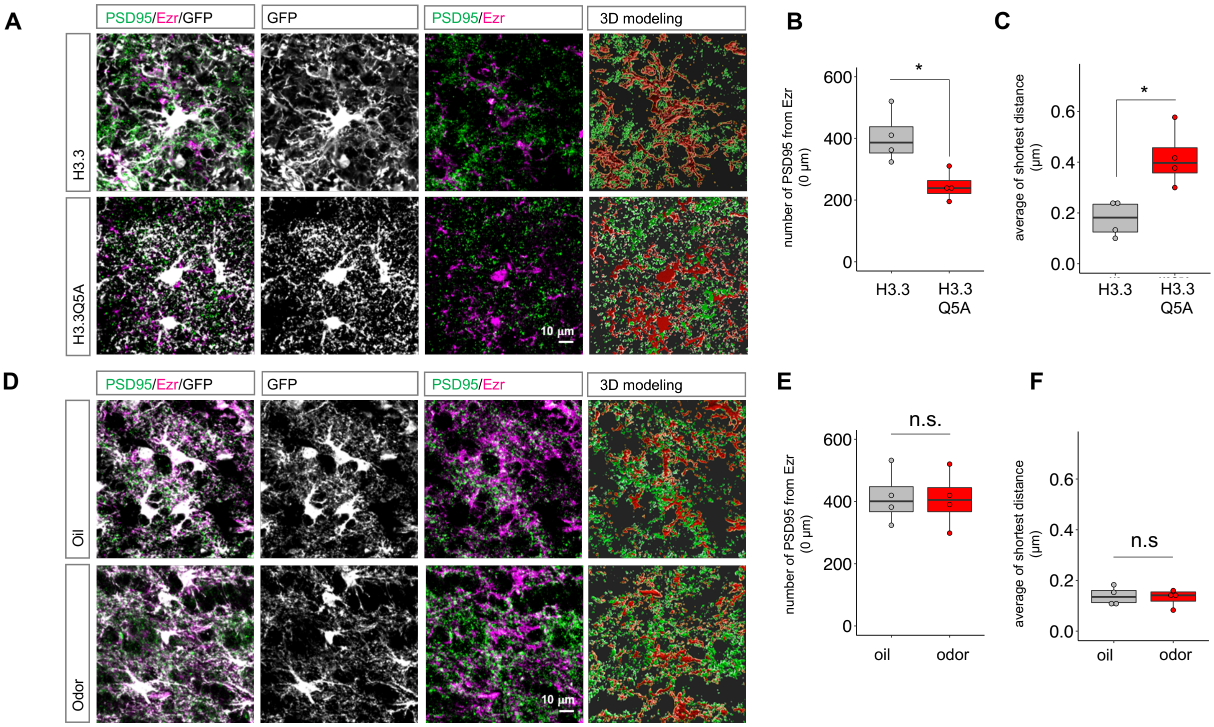


**Supp Fig. S20**

**(A-B)** Representative images and quantification of ezrin and PSD95 at astrocyte terminal process points in H3.3 vs H3.3Q5A (**p*= 0.0178; **p**= 0.0128) and (**E-F**) in odor vs. mineral oil exposed OBs (*p*= 0.9128; *p*= 0.8025). Each data point average of 12 cells across 3 sections; unpaired Student's two-tailed t-test on n= 4 mice/cohort.

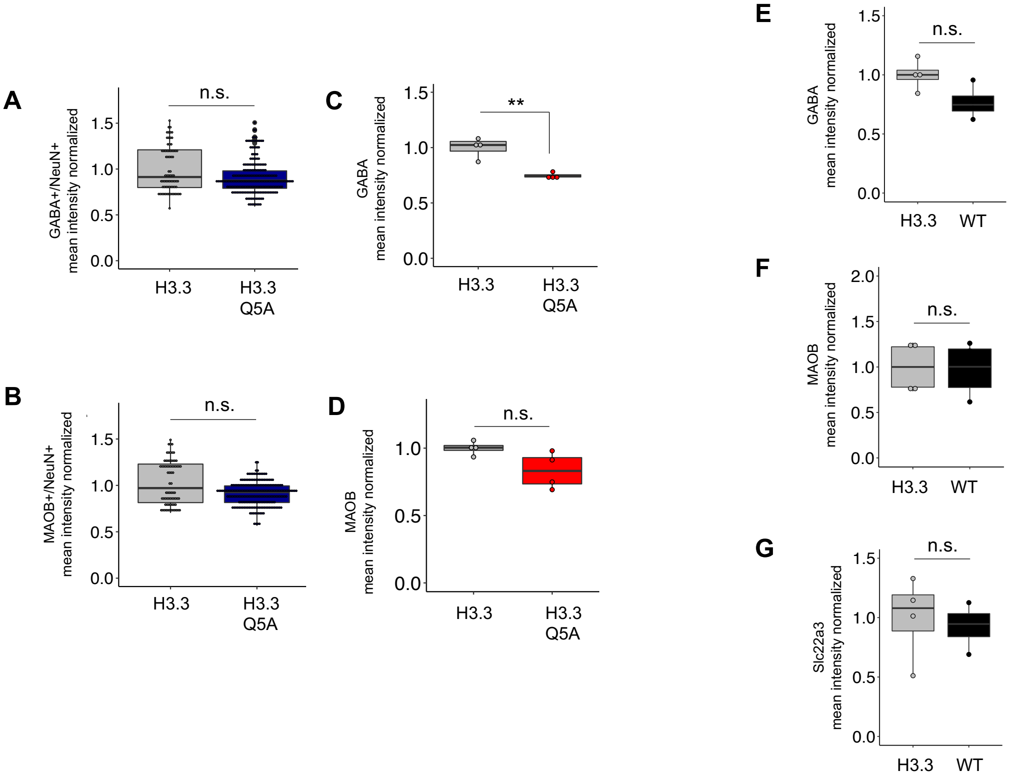


**Supp Fig. S21**

(**A-D**) Quantification of GABA and MAOB in neurons and whole field levels in H3.3Q5A vs. H3.3 control OBs (GABA+/NeuN+: 144 cells/cohort, *p*= 1, Wilcoxon rank sum test on n= 4 mice/cohort; MAOB+/NeuN+: 144 cells/cohort, *p*= 0.2835, unpaired Student's two-tailed t-test on n= 4 mice/cohort; GABA: each data point average of 3 sections, ***p*= 0.0017, unpaired Student's two-tailed t-test on n= 4 mice/cohort; MAOB: each data point average of 3 sections, *p*= 0.0571, Wilcoxon rank sum test on n= 4 mice/cohort). (**E-F**) Quantification of GABA, MAOB and Slc22a3 in H3.3 control vs Aldh1l1-GFP OBs (GABA *p*= 0.0504, MAOB *p*= 0.8889, Slc22a3 *p*= 0.7262, each data point average of 3 sections, unpaired Student's two-tailed t-test on n= 4 mice/cohort).

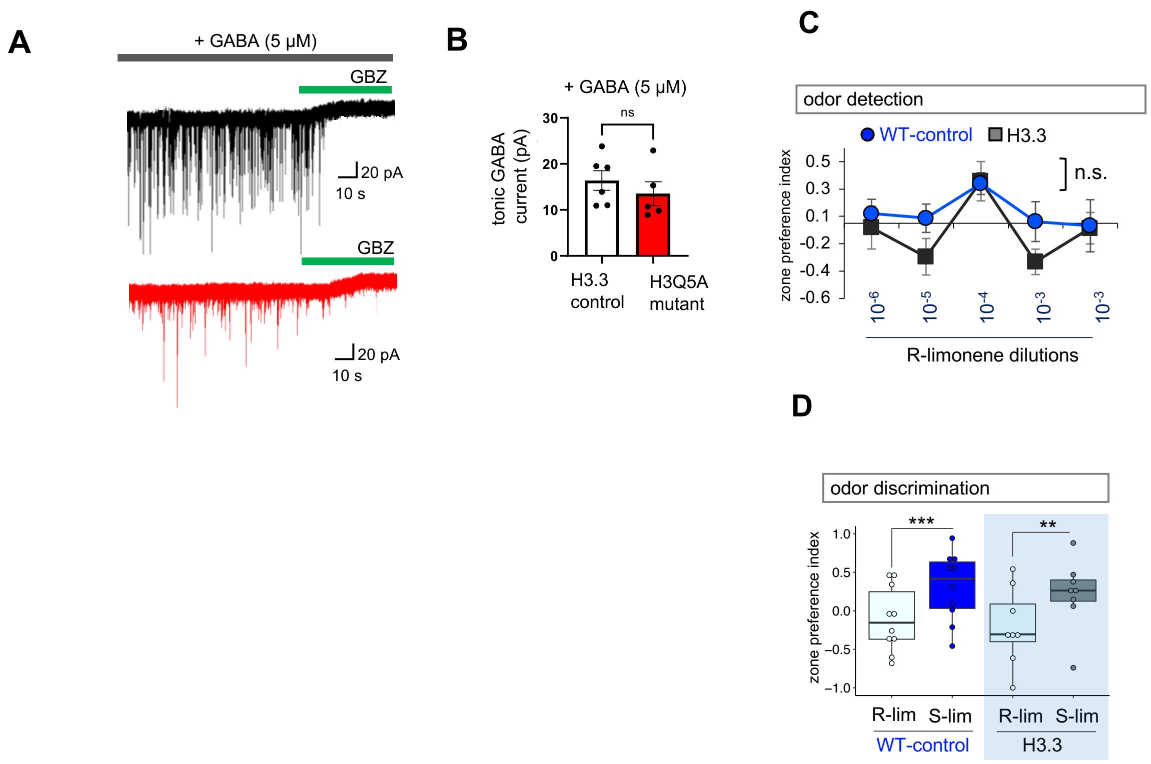


**Supp Fig. S22**
(**A-B**) Representative traces and quantification of measurement of tonic GABA current in presence of GABA (5-6 cells/cohort, *p*= 0.4617, unpaired two-tailed Student's t-test on n= 3 mice/cohort). Data presented as mean ± SEM. GBZ: gabazine, 20 μM. (**C-D**) Quantification of odor detection in H3.3 compared to wild-type mice (n=8-10 each cohort, *p*= 0.6937; two-way repeated measures ANOVA with Sidak multiple comparison). (**J**) Quantification of odor discrimination between R-lim and S-lim from the same cohorts of mice (n=8-10 each cohort, ***p= 0.0008; two-way repeated measures ANOVA with Sidak multiple comparisons).
